# Cortico-subcortical converging organization at rest

**DOI:** 10.1101/2025.01.26.634847

**Authors:** Dheemant Jallepalli, Shilpa Dang

## Abstract

Higher-order cognitive function relies on information convergence between functionally specialized communities. It is unknown how subcortex configures itself with respect to the local segregation and global integration dynamics at rest. Using functional MRI, we revealed three non-overlapping communities confined anatomically to thalamus, basal ganglia, and subcortical limbic structures and network “hubs”. Further, using statistical modelling we quantified the extent of convergence and found multiple, widespread cortical regions converging onto individual subcortical regions. We revealed a topographic organization of cortical convergence within subcortical resting state networks, such that thalamic, basal ganglia, subcortical limbic networks received convergence from cortical areas involved in sensorimotor integration, associative and motor control, limbic functions, respectively. Further, we found functional diversity of cortex to be the major driving factor behind cortical convergence within subcortex and that the absence of subcortical regions significantly impacted the information-transmission efficiency within the converging organization, underscoring the subcortical contributions to brain dynamics.

---

The human subcortex lies deep within the brain and comprises ∼8% of brain mass but contains only 0.8% of neurons in human brain^1^. Despite its size, subcortex consists of diverse neural structures, including thalamus, basal ganglia, hippocampus and amygdala. These structures have been assumed, in past, to simply subserve cortex^2^. Contrary to this perspective, subcortical structures perform specific functions^3–6^ through subcortical and cortico-subcortical circuits^7–9^. These circuits are parallelly organized, functionally segregated, and linked with goal-directed and habitual behaviours^7,10,11^ (**Fig. 1A**). However, higher-order cognitive functions such as goal-directed behaviours cannot occur in isolation, instead they rely on integration of information from various specialized neural communities^9,12–15^. Past studies have shown convergence from functionally diverse cortical sites within the subcortex^16–18^. Moreover, for both functional segregation and integration the subcortex serves as an important neuroanatomical substrate due to its key anatomical features, including progressive compression of pathways over incrementally smaller structures^19^ and proximity of many smaller nuclei^1^ (**Fig. 1A)**. We believe that these features are notably beneficial for information integration, as it allows for synaptic contact and thus, communication between proximal subcortical regions of distinct functional systems^7,10,20^.

**Figure 1.**
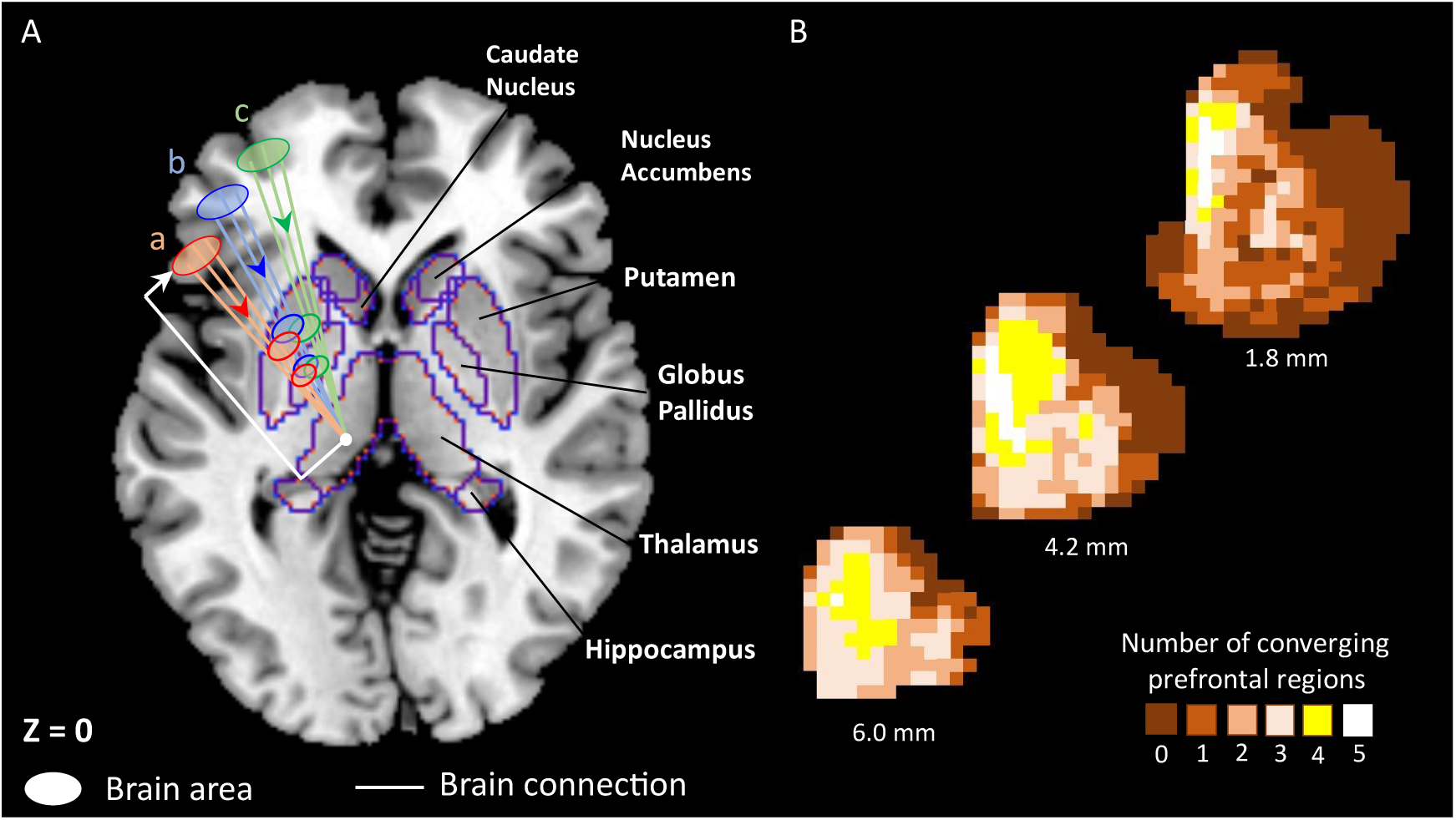
Evidence from past. **A** Generalized schematic diagram based on the parallel organization of functionally segregated circuits^7^. Cortical connections from areas (a, b, c) partially converge in the dorsal striatum (a structure of basal ganglia consisting of putamen and caudate nucleus). The striatal regions send further converging connections to incrementally smaller areas such as globus pallidus, which in turn project to a specific region of the thalamus. The outputs from these basal ganglia structures loop back to the cortex via thalamus in parallel but segregated basal ganglia – thalamocortical channels, specifically named as motor circuit, oculomotor circuit, lateral orbitofrontal circuit, dorsolateral prefrontal circuit, and anterior cingulate circuit^7^. Arrows indicate direction of the brain connection. **B** Heatmaps of cortico-striatal convergence areas in rhesus macaques, based on data from an invasive tract-tracing experiment^17^. These heatmaps represent voxel-wise converging projections in the striatum from 0 to 5 distinct prefrontal cortical (PFC) regions (i.e., ventromedial PFC, orbitofrontal cortex, dorsal PFC, ventrolateral PFC, dorsal anterior cingulate cortex). Shown here are striatal slices at 6.0 mm, 4.2 mm, and 1.8 mm, anterior to anterior commissure (recreated representative slices with permission).

Despite above findings, it is unknown how subcortex configures itself at rest with respect to the local segregation and global integration dynamics^14^. Unlike prevalent research on cortical resting state networks (RSNs)^21,22^, we found that the RSN organization within subcortex (to the best of our knowledge) has not been studied explicitly and the available knowledge is fragmented and with respect to cortex only^23^. We hypothesized that having specialized functional roles would drive the formation of functionally segregated RSNs of subcortical regions. Thus, first we determined the RSNs of subcortex using functional connectivity and clustering algorithm. Additionally, we examined the topology of subcortex and compared it with that of cortex using network science. Further, the evidence for convergence of cortical connections within subcortex in humans at rest is limited and stereotypical. Several functional neuroimaging studies have focused on cortico-subcortical correlations during RS^24–27^, providing evidence that spontaneous neural activity in subcortex can be fractionated into several independent components, each correlating with activity in cortical RSNs. Strikingly, these neuroimaging studies have adopted a cortico-centric approach and focussed on delineating the network-level components within subcortex rather than discovering the intricacies of the organization of cortical connections within the subcortex at region-level. Elucidating the cortico-subcortical convergence at rest is crucial for two overarching goals of the field – one, to understand how efficient brain function is based on successful communication between subcortex and cortex and two, how alterations in this communication may lead to various diseased states^11,28,29^.

Much remains to be known about the cortico-subcortical converging organization at rest in humans. Specifically, we do not know the exact cortical regions converging onto subcortical regions, what causes these cortical regions to converge, and what makes subcortex indispensable for convergence. As a central aim of this study, we examined the organization of cortical connections converging within the subcortex. Using RS functional neuroimaging data of 92 healthy adult participants^30^, statistical modelling, and network science^15^ – first, we computed the extent of convergence from distinct cortical regions within the subcortex. Next, based on past evidence^7,16,17,19^, we examined how convergence between the cortical regions was related to the distance and type of nodes (functionally diverse or similar) in the cortex. Finally, we validated the indispensable role of subcortex by simulating a targeted attack on subcortical nodes and assessing its potential impact on information transmission efficiency of the converging organization.

## Results

### Resting state networks of subcortex

We determined the segregated communities in subcortex at rest using group-level functional connectivity (FC) of 27 subcortical nodes or regions-of-interest (ROIs). The ROIs were formed by combining bilateral parcellations (shown in **Supplementary Fig. S1**). The FC was computed using partial correlation between the nodes^31^. We employed a standard clustering algorithm that identified communities by maximizing modularity, i.e. stronger connections within the community in comparison to connections between the communities. This analysis revealed 3 functionally segregated communities in the subcortex (**Fig. 2)**. Notably, these communities were consistent with the well-known anatomical structures of the subcortex, i.e. the thalamic network (THA) comprising of all thalamus (THA) nodes, the basal ganglia network (BGN) comprising of all putamen (PUT), caudate nucleus (CAU), nucleus accumbens (NAC), and globus pallidus (GPL) nodes, and the subcortical limbic network (SLN) comprising of all hippocampus (HIP) and amygdala (AMY) nodes. Initially, one of the thalamic nodes, the dorsal anterior region of thalamus (THA-DA), was classified as part of BGN due to its stronger connectivity with BG nodes. However, later we reassigned that node to the thalamic network for consistency with contiguous anatomical boundaries^21^.

**Figure 2.**
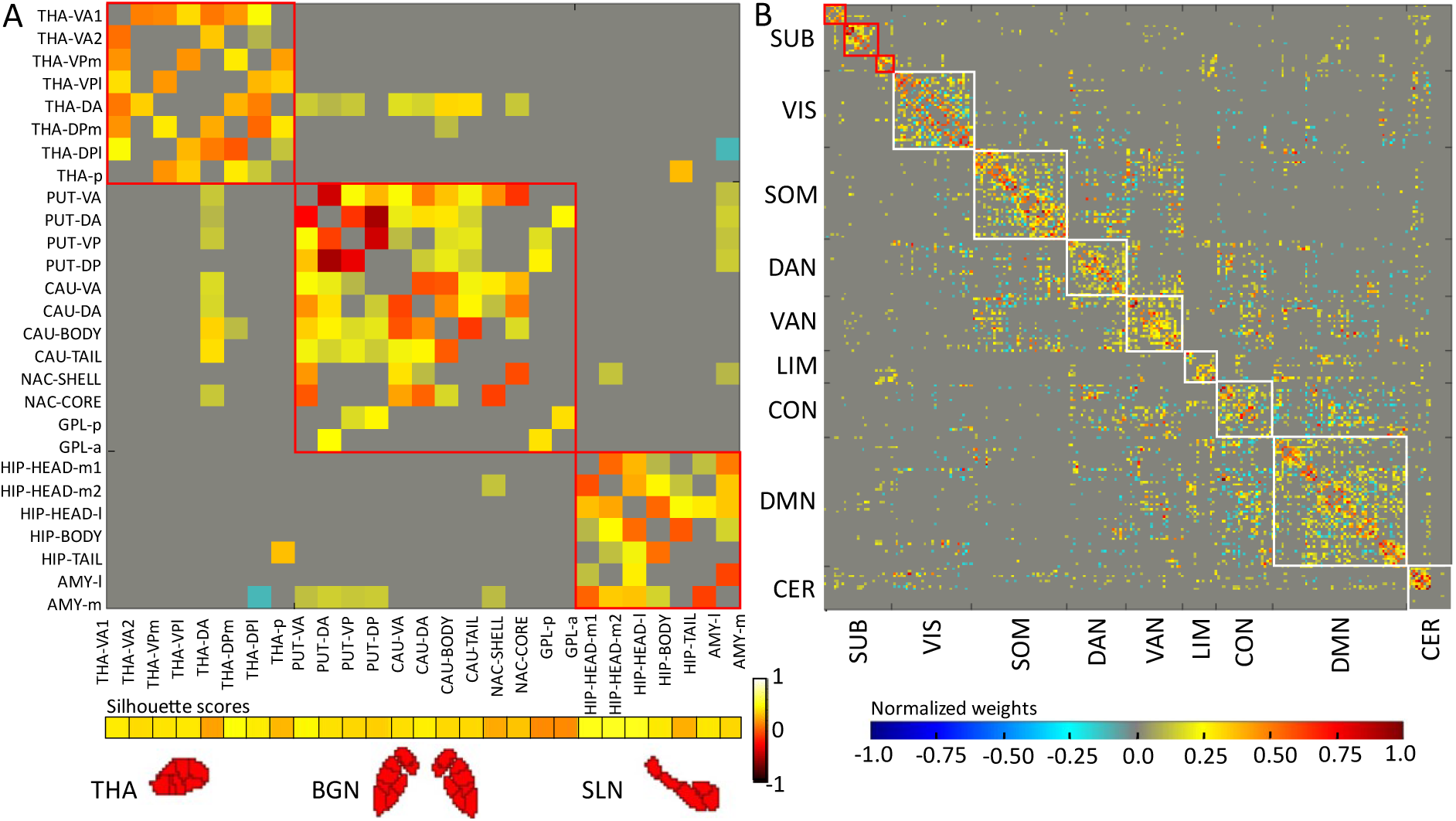
Resting state networks. **A** Group-level ROI-by-ROI weighted functional connectivity matrix based on partial correlation method, sorted by three communities identified within the subcortex, i.e. the thalamic network (THA, comprising of all thalamus ROIs), the basal ganglia network (BGN, comprising of all putamen, caudate nucleus, nucleus accumbens, and globus pallidus ROIs), and the subcortical limbic network (SLN, comprising of all hippocampus and amygdala ROIs), (ROIs: V – ventral, D – dorsal, A or a – anterior, P or p – posterior, l – lateral, m – medial). The block-like pattern along the diagonal indicates a modular organization in subcortex where connections within a community are stronger than connections between the communities. The horizontal bar (at the bottom) represents the node-wise Silhouette scores (ranging from −1.0 𝑡𝑜 1.0) of 27 subcortical ROIs. **B** Similar matrix shown at whole brain level, sorted according to the three newly identified subcortical communities (SUB) and eight existing cortical communities from literature^21,22^ – visual network (VIS), somatomotor network (SOM), dorsal attention network (DAN), ventral attention network (VAN), limbic network (LIM), control network (CON), and default mode network (DMN). The horizontal jet colormap (at the bottom) represents the strength of normalized weights (ranging from −1.0 to 1.0) for both the panels A and B.

**Fig. 2** shows the group-level ROI-by-ROI weighted FC matrix for the subcortex and the whole brain, sorted by community assignment. The block-like patterns in the matrix indicate that the subcortical RS organization exhibited modularity, i.e. stronger partial correlation within-community than between-community (statistics reported in next section along with other metrics). This was similar to the existing RS organization found in cortex^21,22^. Moreover, for comparison we also computed the group-level pair-wise full correlations between subcortical nodes and found no clear block-like pattern for the identified communities (**Supplementary Fig. S2**). This indicated that the partial correlation reliably detected the true connectivity between nodes^31^ and thus, in turn the true modular organization.

For validation, silhouette score (SS) was used. SS indicates the confidence score of a node’s affiliation with the assigned community. We found that the scores for subcortical nodes were all positive and significantly greater than zero (mean SS (± SD) = 0.39 (±0.11), *t*[26] = 18.61, *p* < 0.01, horizontal bar at bottom of **Fig. 2A**), indicating an appropriate clustering solution. Further, we measured the topological characteristics using node-level and network-level characteristic graph metrics of segregation and integration^15^. Statistics-based comparison showed that the topological characteristics of the subcortex were significantly different than expected in a “null” distribution obtained from a population of 1000 random topologies^32,33^ (**Fig. 3** and **Supplementary Table S1**). Overall, these results validated the reliability of the identified modular organization in the subcortex.

**Figure 3.**
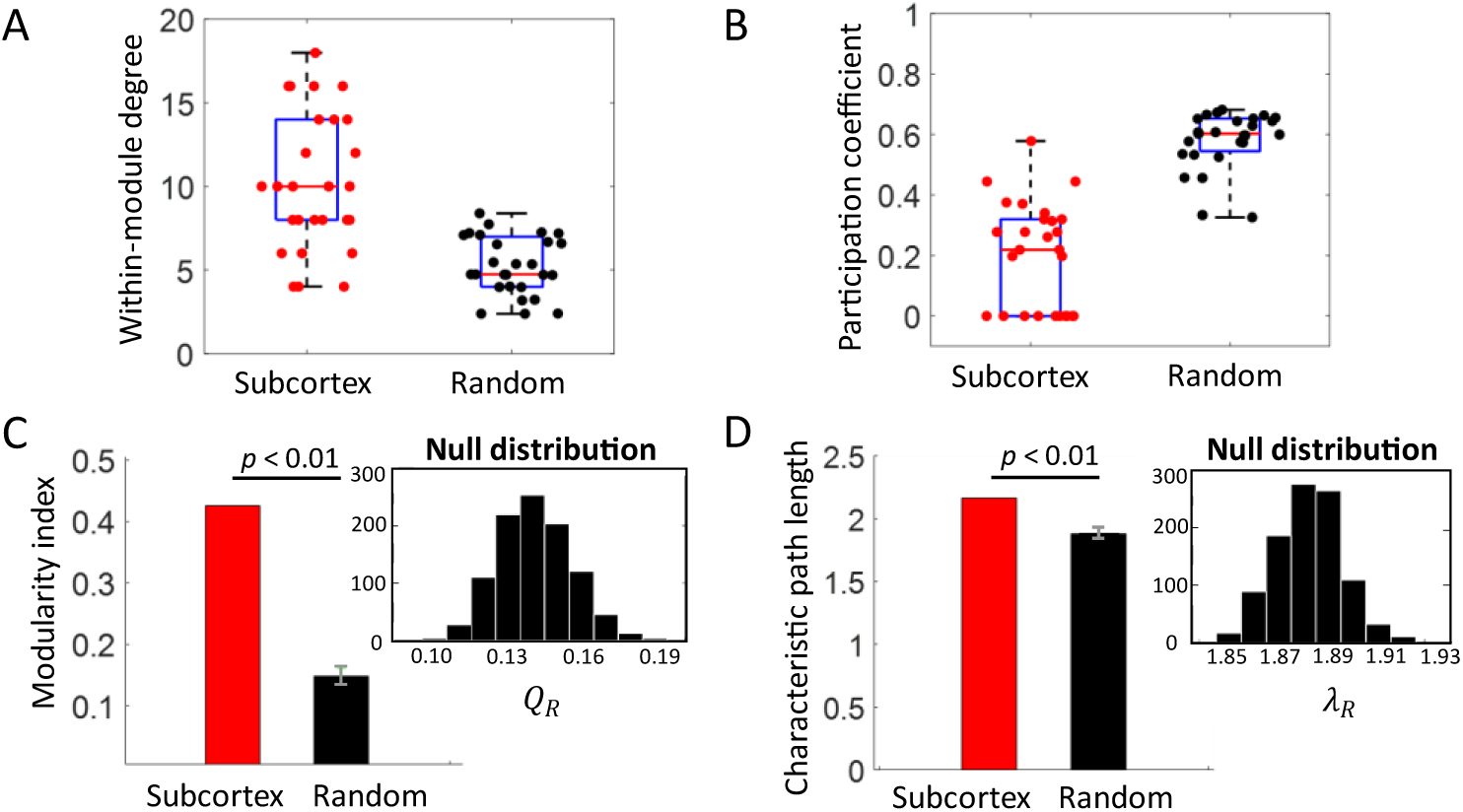
Characteristic graph metric values across subcortex and random networks. **A-B** Node-level metrics. Higher within-module degree (WMD) and lower participation coefficient (PC) of 27 subcortical ROIs (red dots) than their counterpart nodes in random network (black dots), illustrating significant difference in the topology of subcortex in comparison to random (R) network (mean *wmd*_*i*_ (± SD) = 10.59 (±3.85), mean *wmd*_*iR*_ (± SD) = 5.66 (±1.84), mean 𝑃_*i*_ (± SD) = 0.20 (±0.18), mean 𝑃_*iR*_ (± SD) = 0.58 (±0.09); all corr. *p*-values < 0.01, *i* = 1 𝑡𝑜 27, see **Supplementary Table S1** for node-wise statistics). Please note that the PCs have been calculated with respect to subcortical communities only. Each red dot represents metric value of one node in the subcortex and each black dot represents the mean metric value of the “null” distribution obtained from a population of 1000 random networks. On each box, the central mark (solid red line) is the median, the edges of the box are the 25^th^ and 75^th^ percentiles, the whiskers extend to the maximum and minimum datapoints. **C-D** Network level metrics. Higher modularity index 𝑄 and characteristic path length 𝜆 of subcortex network (red bars) than random network (black bars), illustrating stronger modular organization in subcortex than expected in random networks (𝑄 = 0.43, mean 𝑄_*R*_ (± SD) = 0.15 (±0.02); *t*[999] = 686.58, corr. *p* < 0.01; 𝜆 = 2.16, mean 𝜆_*R*_ (± SD) = 1.88 (±0.01); *t*[999] = 737.66, corr. *p* < 0.01). Black bar in each main panel represents mean metric value of the “null” distribution obtained from a population of 1000 random networks (see inset panel). Error bars (in grey) indicate standard deviation of metric values across random networks.

### Subcortical hubs

We examined the topology of subcortex to understand its role in the whole brain organization at rest. For this, we identified four distinct types of nodes, including connector hubs, non-hub connectors, provincial hubs, and non-hubs, based on a fundamental work^34^. Hubs are brain regions with high degree that play a key role in information processing for functional segregation and integration^15,33^. We found that about 81.48% of nodes in subcortex (27 nodes) were connectors (15 connector hubs and 7 non-hub connectors) and remaining 18.52% were non-connector type (1 provincial hub and 4 non-hubs) (**Fig. 4A-C**). Further, we found a similar distribution in cortex (217 nodes), as 81.11%:18.89%. Moreover, we found that the topological characteristics, including connection density (degree per voxel), clustering, and participation coefficients were comparable across subcortex and cortex (**Fig. 4D-I** and **Supplementary Table S2**).

**Figure 4.**
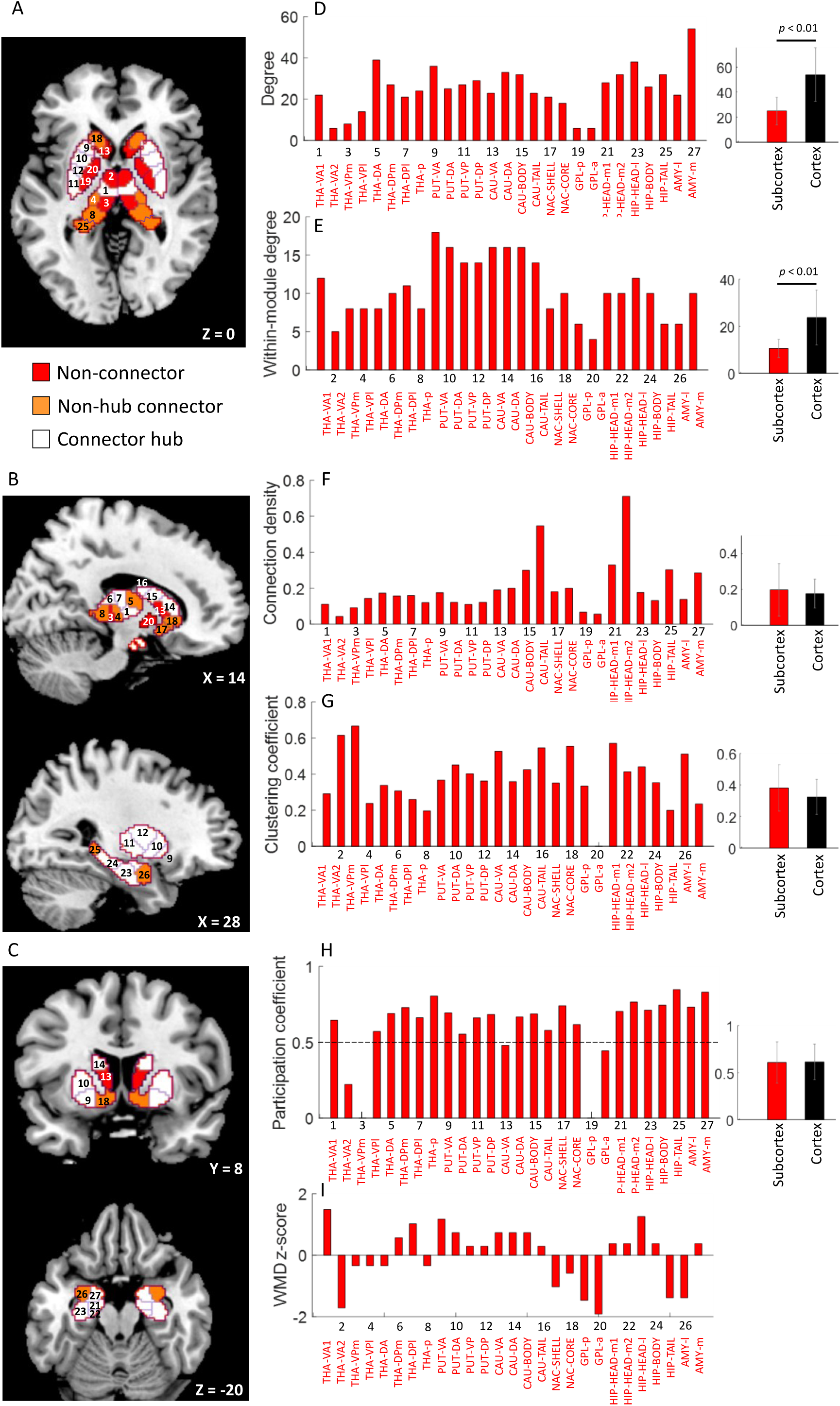
Topology of subcortex and graph metrics. **A-C** Distinct types of nodes in the subcortex crucial for balancing the local segregation and global integration dynamics – the non-connectors (in red) including 4 non-hubs (THA-VA2, THA-VPm, GPL-p, GPL-a) and 1 provincial hub (CAU-VA), the latter type are important for specialised functional computations, 7 non-hub connectors for global information integration only (in orange; THA-VPl, THA-DA, THA-p, NAC-shell, NAC-core, HIP-tail, AMY-l), and 15 connector hubs for both functional segregation and integration (in white; THA-VA1, THA-DPm, THA-DPl, PUT-VA, PUT-DA, PUT-VP, PUT-DP, CAU-DA, CAU-body, CAU-tail, HIP-head-m1, HIP-head-m2, HIP-head-l, HIP-body, AMY-m). Numbers 1 − 27 are shown node-wise for reference. **D-I** Node-wise values of (within-module) degree, connection density, clustering coefficient, participation coefficient, and within-module degree z-score of each of the 27 nodes in the subcortex. Also shown in right-most panels is the comparison across the topology of subcortex (27 nodes) and cortex (217 nodes). The bars in rightmost panels of D-I represent mean values across respective nodes. Error bars (in grey) indicate standard deviation of metric values across nodes. Significant difference across graph metrics of subcortex and cortex was determined using a two-sample t-test with equal means and equal but unknown variances at *p* < 0.01 (see **Supplementary Table S2** for individual graph metrics’ statistics). Please note that the PCs have been calculated with respect to whole brain communities.

### Cortico-subcortical converging organization at rest

#### Extent of convergence within subcortex

We quantified the cortical convergence within subcortex by computing the absolute number of cortical ROIs converging onto each of the 27 subcortical ROIs. This analysis not only revealed information about how many cortical regions converged but also specifically where that integration of information occurred and among which of the cortical ROIs (**Fig. 5** and **Table 1**). The absolute number of cortical (bidirectional) connections onto a subcortical ROI ranged from 0 − 16, where non-connectors received at most from one cortical site and connectors received from multiple cortical sites (**Fig. 5A**). Notably, we observed that the convergence within subcortex was topographically organized with respect to the large-scale network organization at rest^21,22^, presented next.

**Figure 5.**
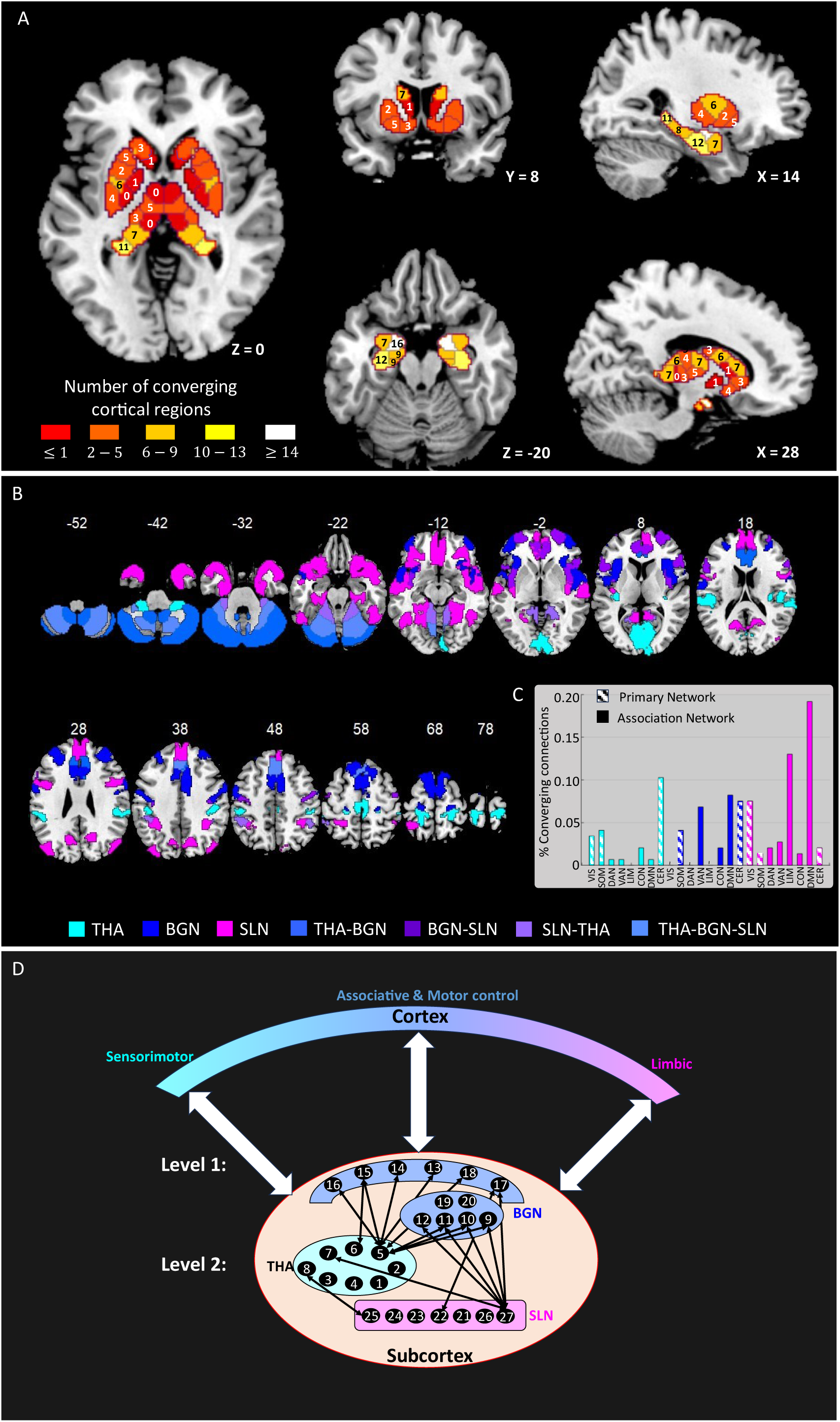
Cortico-subcortical converging organization at rest. **A** Areas of convergence in the human subcortex. Each sub-panel shows the number of distinct cortical regions converging (via bidirectional connections) onto each of the 27 regions-of-interest within the subcortex. Colour and number on each subcortical ROI indicate the range of convergence and the exact number of converging inputs, respectively. **B** Cortical regions converging onto subcortical ROIs at a significance threshold of *p <* 0.01, Bonferroni corrected for multiple comparisons. For better visualization shown separately for ROIs of thalamic network (THA, in cyan), basal ganglia network (BGN, in blue), and subcortical limbic network (SLN, in magenta). Some cortical regions converge to ROIs of more than one subcortical network and are color-coded distinctly (shades of blue and purple mentioned at the bottom of panel B). Axial slices shown at different z-coordinates. **C** Percentage of total number of converging (bidirectional) connections between a cortical and a subcortical resting state network. The % converging connections on subcortical networks THA, BGN, SLN are shown using distinct coloured bars (cyan, blue, magenta, respectively), including those from primary, i.e. early sensory and late motor related networks including cerebellum as patterned bars whereas from association networks as solid bars. **D** Topographic organization of functionally segregated cortico-subcortical circuits existing at rest and their integration within subcortex. Level 1 represents convergence from functionally diverse cortical regions onto subcortex, which is broadly separated according to functions – sensorimotor integration, associative, motor control, and limbic and shows mapping onto distinct subcortical networks THA, BGN, and SLN. Level 2 represents further convergence between functionally segregated cortico-subcortical circuits via between-community interactions within subcortex (see Fig. 2 for reference; within-community connections are not included here). Numbers 1 − 27 represent the 27 subcortical nodes/ROIs. Bidirectional arrows indicate brain connection.

**Table 1.** Converging cortical regions within subcortex.

| Table 1. Converging cortical regions within subcortex |  |  |  |  |  |  |
| --- | --- | --- | --- | --- | --- | --- |
| ROI index | Subcortical ROI | Number of converging connections | Converging cortical ROIs* | Centroids (x,y,z) (in mm) of bilateral parcellations forming a cortical ROI | RSN | T(91)** |
| 1 | THA-VA1 | 5 | V1 | (-5,-88,2), (8,-92,-2) | VIS | 4.29 |
|  |  |  | Area 4 (M1) | (-4,-25,56), (4,-25,58) | SOM | 4.32 |
|  |  |  | Area p24 (PreACC) | (-5,34,20), (7,35,25) | DMN | 4.54 |
|  |  |  | CER 6 | (-22,-61,-24), (26,-60,-25) | CER | 5.80 |
|  |  |  | CER 9 | (-10,-51,-48), (11,-51,-48) | CER | 7.65 |
| 2 | THA-VA2 | 0 | - | - | - | - |
| 3 | THA-VPm | 0 | - | - | - | - |
| 4 | THA-VPl | 3 | OP1 (S2) | (-48,-24,18), (49,-21,18) | SOM | 5.72 |
|  |  |  | Area 3b (S1) | (-18,-31,69), (22,-29,68) | SOM | 4.98 |
|  |  |  | Area 2 (S1) | (-46,-29,44), (45,-28,43) | DAN | 5.08 |
| 5 | THA-DA | 7 | Area 8BM (dmPFC) | (-4,28,47), (5,28,48) | CON | 4.46 |
|  |  |  | CER Crus 1 | (-35,-68,-31), (39,-69,-31) | CER | 5.62 |
|  |  |  | CER Crus 2 | (-27,-75,-40), (33,-71,-42) | CER | 4.58 |
|  |  |  | CER 4, 5 | (-14,-45,-19), (18,-45,-20) | CER | 6.38 |
|  |  |  | CER 6 | (-22,-61,-24), (26,-60,-25) | CER | 4.14 |
|  |  |  | CER 8 | (-25,-56,-50), (26,-58,-51) | CER | 9.03 |
|  |  |  | CER 10 | (27,-36,-43), (-21,-36,-44) | CER | 4.52 |
| 6 | THA-DPm | 6 | Area 3b (S1) | (-18,-31,69), (22,-29,68) | SOM | 4.27 |
|  |  |  | Area 8BM (dmPFC) | (-4,28,47), (5,28,48) | CON | 5.29 |
|  |  |  | CER Crus 1 | (-35,-68,-31), (39,-69,-31) | CER | 4.54 |
|  |  |  | CER 8 | (-25,-56,-50), (26,-58,-51) | CER | 4.58 |
|  |  |  | CER 9 | (-10,-51,-48), (11,-51,-48) | CER | 5.46 |
|  |  |  | CER 10 | (27,-36,-43), (-21,-36,-44) | CER | 5.21 |
| 7 | THA-DPl | 4 | V1 | (-7,-74,10), (9,-73,8) | VIS | 4.62 |
|  |  |  | V1 | (-5,-88,2), (8,-92,-2) | VIS | 4.72 |
|  |  |  | A1 | (-36,-24,10), (35,-22,14) | SOM | 4.47 |
|  |  |  | CER 6 | (-22,-61,-24), (26,-60,-25) | CER | 4.14 |
| 8 | THA-p | 7 | V2 | (-13,-43,-5), (18,-45,-3) | VIS | 8.07 |
|  |  |  | POS1 | (-14,-49,4), (14,-47,4) | VIS | 8.16 |
|  |  |  | Area 2 (S1) | (-19,-40,72), (22,-35,70) | SOM | 4.71 |
|  |  |  | PF (IPL) | (-55,-32,22), (58,-31,24) | VAN | 5.03 |
|  |  |  | PF (IPL) | (-45,-41,47), (47,-44,47) | CON | 5.50 |
|  |  |  | CER 4, 5 | (-14,-45,-19), (18,-45,-20) | CER | 4.41 |
|  |  |  | CER 9 | (-10,-51,-48), (11,-51,-48) | CER | 7.41 |
| 9 | PUT-VA | 5 | FOP | (-36,4,11), (38,7,10) | VAN | 4.33 |
|  |  |  | Area a24pr (MCC) | (-6,22,31), (7,19,35) | VAN | 5.83 |
|  |  |  | Area a47r (vlPFC) | (-42,49,-7), (42,51,-6) | CON | -4.07 |
|  |  |  | Area 9m (dmPFC) | (-6,45,6), (8,42,4) | DMN | 4.96 |
|  |  |  | Area p24 (PreACC) | (-5,34,20), (7,35,25) | DMN | 4.95 |
| 10 | PUT-DA | 2 | Area 4 (M1) | (-59,-2,24), (61,6,29) | SOM | 4.86 |
|  |  |  | SCEF (PMC) | (-5,9,48), (8,2,43) | VAN | 4.97 |
| 11 | PUT-VP | 4 | Area 55b (PMC) | (-51,-7,43), (52,-6,37) | SOM | 4.55 |
|  |  |  | Area 4 (M1) | (-41,-14,48), (44,-11,48) | SOM | 9.75 |
|  |  |  | Area 4 (M1) | (-19,-24,66), (21,-24,66) | SOM | 5.60 |
|  |  |  | CER 6 | (-22,-61,-24), (26,-60,-25) | CER | 7.27 |
| 12 | PUT-DP | 6 | Area 4 (M1) | (-59,-2,24), (61,6,29) | SOM | 4.84 |
|  |  |  | Area 4 (M1) | (-41,-14,48), (44,-11,48) | SOM | 5.41 |
|  |  |  | PoI | (-39,2,-5), (41,9,-3) | VAN | 4.93 |
|  |  |  | FOP | (-50,1,4), (49,5,4) | VAN | 4.14 |
|  |  |  | Area 6ma (SMA) | (-8,-3,71), (7,-2,67) | VAN | 6.02 |
|  |  |  | CER 6 | (-22,-61,-24), (26,-60,-25) | CER | 8.21 |
| 13 | CAU-VA | 1 | FOP | (-33,19,8), (37,23,5) | VAN | 5.09 |
| 14 | CAU-DA | 7 | SCEF (SMA) | (-5,9,48), (8,2,43) | VAN | 4.39 |
|  |  |  | Area a10p (dlPFC) | (-28,58,-1), (27,59,3) | DMN | 5.49 |
|  |  |  | Area p24 (PreACC) | (-5,34,20), (7,35,25) | DMN | 5.52 |
|  |  |  | SFL (SMA) | (-13,24,61), (12,20,63) | DMN | 4.18 |
|  |  |  | SFL (SMA) | (-7,10,64), (6,11,57) | DMN | 6.26 |
|  |  |  | CER Crus 1 | (-35,-68,-31), (39,-69,-31) | CER | 4.13 |
|  |  |  | CER Crus 2 | (-27,-75,-40), (33,-71,-42) | CER | 5.54 |
| 15 | CAU-Body | 6 | AVI | (-33,25,-1), (37,23,5) | VAN | 4.37 |
|  |  |  | Area 8BM (dmPFC) | (-4,28,47), (5,28,48) | CON | 5.17 |
|  |  |  | Area 44 (IFG) | (-53,19,11), (53,24,6) | DMN | 4.54 |
|  |  |  | CER 6 | (-22,-61,-24), (26,-60,-25) | CER | 5.10 |
|  |  |  | CER Crus 1 | (-35,-68,-31), (39,-69,-31) | CER | 5.91 |
|  |  |  | CER Crus 2 | (-27,-75,-40), (33,-71,-42) | CER | 5.94 |
| 16 | CAU-Tail | 3 | STGa | (-53,6,-12), (53,3,-6) | DMN | -4.44 |
|  |  |  | CER 6 | (-22,-61,-24), (26,-60,-25) | CER | 4.36 |
|  |  |  | CER Crus 1 | (-35,-68,-31), (39,-69,-31) | CER | 5.87 |
| 17 | NAC-Shell | 4 | AVI | (-33,16,-8), (34,21,-8) | CON | 4.9 |
|  |  |  | Area 9m (dmPFC) | (-6,45,6), (8,42,4) | DMN | 10.01 |
|  |  |  | Area p24 (PreACC) | (-5,34,20), (7,35,25) | DMN | 5.13 |
|  |  |  | CER 9 | (-10,-51,-48), (11,-51,-48) | CER | 4.91 |
| 18 | NAC-Core | 3 | Area 9/46d (dlPFC) | (-29,43,30), (32,45,28) | VAN | 4.28 |
|  |  |  | Area 9m (dmPFC) | (-6,45,6), (8,42,4) | DMN | 7.5 |
|  |  |  | Area p24 (PreACC) | (-5,34,20), (7,35,25) | DMN | 6.13 |
| 19 | GPL-p | 0 | - | - | - | - |
| 20 | GPL-a | 1 | CER 8 | (-25,-56,-50), (26,-58,-51) | CER | 4.40 |
| 21 | HIP-Head-m1 | 9 | PHC | (-30,-33,-18), (31,-31,-18) | VIS | 4.64 |
|  |  |  | POS1 (VC) | (-14,-49,4), (14,-47,4) | VIS | 7.88 |
|  |  |  | PerC | (-26,-9,-33), (28,-1,-40) | LIM | 6.28 |
|  |  |  | EntC | (-21,-21,-26), (22,-18,-27) | LIM | 9.74 |
|  |  |  | STSda | (-57,-9,-14), (55,-4,-14) | DMN | 5.03 |
|  |  |  | PGp (IPL) | (-40,-79,30), (45,-75,31) | DMN | 4.58 |
|  |  |  | Area 10r (vmPFC) | (-6,35,-9), (5,40,-10) | DMN | 6.10 |
|  |  |  | POS1 (pCunPCC) | (-8,-52,9), (6,-52,23) | DMN | 5.08 |
|  |  |  | POS1 (pCunPCC) | (-13,-61,19), (13,-55,16) | DMN | 7.93 |
| 22 | HIP-Head-m2 | 9 | PHC | (-19,-37,-12), (18,-36,-13) | VIS | 4.11 |
|  |  |  | POS1 (VC) | (-14,-49,4), (14,-47,4) | VIS | 6.28 |
|  |  |  | TGd | (-32,12,-30), (29,12,-31) | LIM | 4.98 |
|  |  |  | EntC | (-21,-21,-26), (22,-18,-27) | LIM | 15.88 |
|  |  |  | STSda | (-57,-9,-14), (55,-4,-14) | DMN | 4.38 |
|  |  |  | PGp (IPL) | (-40,-79,30), (45,-75,31) | DMN | 4.62 |
|  |  |  | Area 10r (vmPFC) | (-6,35,-9), (5,40,-10) | DMN | 5.23 |
|  |  |  | POS1 (pCunPCC) | (-13,-61,19), (13,-55,16) | DMN | 6.78 |
|  |  |  | CER 6 | (-22,-61,-24), (26,-60,-25) | CER | 4.68 |
| 23 | HIP-Head-l | 12 | PHC | (-30,-33,-18), (31,-31,-18) | VIS | 12.01 |
|  |  |  | PerC | (-26,-9,-33), (28,-1,-40) | LIM | 5.82 |
|  |  |  | EntC | (-21,-21,-26), (22,-18,-27) | LIM | 5.64 |
|  |  |  | TGd | (-43,13,-33), (45,14,-30) | LIM | 5.56 |
|  |  |  | TGd | (-32,12,-30), (29,12,-31) | LIM | 6.98 |
|  |  |  | STSda | (-57,-9,-14), (55,-4,-14) | DMN | 6.38 |
|  |  |  | Area a47r (vlPFC) | (-36,37,-13), (35,38,-14) | DMN | 5.68 |
|  |  |  | Area 10v (vmPFC) | (-5,55,-10), (5,40,-10) | DMN | 5.26 |
|  |  |  | Area 10r (vmPFC) | (-6,35,-9), (5,40,-10) | DMN | 4.93 |
|  |  |  | Area 9m (dmPFC) | (-6,45,6), (8,42,4) | DMN | 5.23 |
|  |  |  | POS1 (pCunPCC) | (-8,-52,9), (6,-52,23) | DMN | 4.10 |
|  |  |  | POS1 (pCunPCC) | (-13,-61,19), (13,-55,16) | DMN | 7.22 |
| 24 | HIP-Body | 8 | PHC | (-19,-37,-12), (18,-36,-13) | VIS | 13.29 |
|  |  |  | PHC | (-30,-33,-18), (31,-31,-18) | VIS | 6.63 |
|  |  |  | TGd | (-43,13,-33), (45,14,-30) | LIM | 6.19 |
|  |  |  | TGd | (-32,12,-30), (29,12,-31) | LIM | 5.74 |
|  |  |  | Area 47s (vlPFC) | (-36,22,-15), (35,23,-17) | DMN | 4.34 |
|  |  |  | Area a10p (dlPFC) | (-28,58,-1), (27,59,3) | DMN | -5.05 |
|  |  |  | Area 9m (dmPFC) | (-6,45,6), (8,42,4) | DMN | 4.45 |
|  |  |  | CER 4, 5 | (-14,-45,-19), (18,-45,-20) | CER | 6.10 |
| 25 | HIP-Tail | 11 | PHC | (-30,-33,-18), (31,-31,-18) | VIS | 4.13 |
|  |  |  | VMV1 | (-25,-54,-8), (26,-52,-9) | VIS | 7.02 |
|  |  |  | V2 | (-13,-43,-5), (18,-45,-3) | VIS | 6.05 |
|  |  |  | POS1 (VC) | (-14,-49,4), (14,-47,4) | VIS | 10.65 |
|  |  |  | Area 2 (S1) | (-30,-46,63), (31,-41,64) | SOM | 5.03 |
|  |  |  | AIP | (-33,-46,41), (36,-45,45) | DAN | 4.21 |
|  |  |  | PoI | (-39,2,-5), (41,9,-3) | VAN | 4.17 |
|  |  |  | PerC | (-26,-9,-33), (28,-1,-40) | LIM | 4.30 |
|  |  |  | PF (IPL) | (-45,-41,47), (47,-44,47) | CON | 5.46 |
|  |  |  | POS2 (pCunPCC) | (-10,-70,31), (13,-71,39) | DMN | 5.43 |
|  |  |  | CER 8 | (-25,-56,-50), (26,-58,-51) | CER | 4.47 |
| 26 | AMY-l | 7 | STGa | (-44,5,-16), (41,5,-16) | VAN | 4.69 |
|  |  |  | PoI | (-40,-14,-2), (40,-11,-4) | VAN | 4.48 |
|  |  |  | PerC | (-26,-9,-33), (28,-1,-40) | LIM | 4.34 |
|  |  |  | TGd | (-25,6,-39), (28,-1,-40) | LIM | 4.09 |
|  |  |  | TGd | (-43,13,-33), (45,14,-30) | LIM | 8.41 |
|  |  |  | TGd | (-32,12,-30), (29,12,-31) | LIM | 11.26 |
|  |  |  | STSda | (-57,-9,-14), (55,-4,-14) | DMN | 4.36 |
| 27 | AMY-m | 16 | Area 4 (M1) | (-41,-14,48), (44,-11,48) | SOM | 4.43 |
|  |  |  | PH (TPOJ) | (-49,-56,-16), (50,-49,-18) | DAN | 4.57 |
|  |  |  | Area 6r (SMA) | (-49,6,26), (49,8,25) | DAN | 6.17 |
|  |  |  | STGa | (-44,5,-16), (41,5,-16) | VAN | 6.05 |
|  |  |  | Area 13l (OFC) | (-24,22,-20), (23,22,-20) | LIM | 5.21 |
|  |  |  | PerC | (-26,-9,-33), (28,-1,-40) | LIM | 8.41 |
|  |  |  | TGd | (-43,13,-33), (45,14,-30) | LIM | 7.32 |
|  |  |  | TGd | (-32,12,-30), (29,12,-31) | LIM | 11.16 |
|  |  |  | Area 8BM (dmPFC) | (-4,28,47), (5,28,48) | CON | -4.83 |
|  |  |  | STSda | (-57,-9,-14), (55,-4,-14) | DMN | 6.52 |
|  |  |  | A5 (STG) | (-61,-13,-3), (62,-19,0) | DMN | 4.81 |
|  |  |  | Area 47s (vlPFC) | (-36,22,-15), (35,23,-17) | DMN | 5.42 |
|  |  |  | Area a47r (vlPFC) | (-36,37,-13), (35,38,-14) | DMN | 5.27 |
|  |  |  | Area 10r (vmPFC) | (-6,35,-9), (5,40,-10) | DMN | 5.20 |
|  |  |  | Area 44 (IFC) | (-53,19,11), (53,24,6) | DMN | -5.42 |
|  |  |  | Area 9m (dmPFC) | (-4,51,28), (6,52,27) | DMN | 4.34 |

\*ROI names from Glasser atlas <sup>42</sup>; see **Supplementary Table S3** for individual functional roles and references; a – anterior, d – dorsal, l – lateral, m – medial, p – posterior, pr – prime, r – rostral, s – superior, v – ventral, A1 – Primary Auditory Cortex, A5 – Fifth Auditory Area, AIP – Anterior Intra Parietal, AVI – Anterior Ventral Insula, CER – Cerebellum, EntC – Entorhinal Cortex, FC – Frontal Cortex, FOP – Frontal Opercular, IFG – Inferior Frontal Gyrus, INS – Insula, IPL – Inferior Parietal Lobule, M1 – Primary Motor Cortex, MCC – Middle Cingulate Cortex, OFC – Orbitofrontal Cortex, OP – Parietal Operculum, PF – Parietal area F, PG – Parietal area G, pCunPCC – Precuneus-Posterior Cingulate Cortex, PerC – Perirhinal Cortex, PFC – Prefrontal Cortex, PHC – Parahippocampal Cortex, PMC – Premotor Cortex, PoI – Posterior Insula, POS – Parieto-Occipital Sulcus, PreACC – Pregenuar Anterior Cingulate Cortex, S1 – Primary Somatosensory Cortex, S2 – Secondary Somatosensory Cortex, SCEF – Supplementary and Cingulate Eye Fields, SFL – Superior Frontal Language Area, SMA – Supplementary Motor Area, STG – Superior Temporal Gyrus, STS – Superior Temporal Sulcus, TPOJ – Temporo-Parieto-Occipital Junction, TG – Temporal Gyrus, V1 – Primary Visual Cortex, V2 – Second Visual Area, VC – Visual Cortex, VMV – Ventromedial Visual Area
\*\*Individual T-value T(91) reported for a cortical ROI having significant FC with a subcortical ROI computed at group-level (92 participants, random effects analysis (RFX) one-sample *t*-test at corr. *p* < 0.01, see Methods on Functional connectivity)

#### Topographic organization of convergence within subcortex

During resting state, it has been shown that the early sensory and late motor cortices, i.e. visual and somatomotor networks, are separated from the association networks, including the dorsal attention, ventral attention, limbic, control, default mode networks^21^. We found that majority of cortical regions converging onto a particular subcortical network were either from primary or association networks (**Table 1** and **Fig. 5B-C**). The thalamic network had major convergence (about 81% of its connections) from visual, somatomotor networks, and cerebellum. Interestingly, the maximum connections to thalamus were received from cerebellum, which is crucial for locomotion and balance^35^ as well as generation of appropriately timed motor response to sensory stimuli^36^. The remaining were from association networks DAN, VAN, CON, and DMN – particularly, from regions involved in interoception and action inhibition (Area p24), somatosensory processes (OP1), motor planning and execution of goal-directed behaviours, action observation and imitation (Areas 8BM, PF), (**Supplementary Table S3** includes detailed functional roles of converging regions). This indicates the primary involvement of thalamic network in integration of (multimodal) sensory and motor information from cortex^4^(**Fig. 5D**).

Further, the basal ganglia network had major convergence (about 60% of its connections) from widespread frontal, insular, cingulate regions (of VAN, CON, DMN) and remaining from motor cortex and cerebellum. We found evidence that cortico-basal ganglia circuits were involved in planning, learning, and controlling of speech^37^ and other actions. There were converging connections from areas involved in speech production, retrieval, and learning (frontal areas FOP, SFL, 6ma, 44, a47r, 55b), working memory and cognitive control (frontal areas 9m, a10p, 9-46d), risky decision making, socioemotional processing, interoception and autonomic control, attention and saliency processing (AVI), skeletomotor regulation, action selection and inhibition (cingulate areas a24pr, p24), (oculo)motor planning and control in goal-directed behaviours (motor areas SCEF, 4, 6ma, 8BM) (**Supplementary Table S3**). Strikingly, no direct connections were received from the early sensory, dorsal attention, and limbic cortices. However, we believe that any sensory input required for control and refinement of actions^38^ enters these circuits via two pathways – subcortico-subcortical connectivity between thalamus (THA-DA) and BGN nodes (**Fig. 2A**) and converging connections from sensory association cortical areas (PoI, STGa). Similarly, through connectivity with subcortical limbic structures, the BGN nodes provide a gateway for limbic drives to gain access and bias the cortico-basal ganglia circuits^11,39^. Together, this points toward the presence of associative and motor control cortico-basal ganglia circuits, which control and develop non-innate actions/action-sequences during goal-directed and habitual behaviours^11,40^(**Fig. 5D**). The former circuits involve convergence from association regions and latter from movement-related regions^11^.

Finally, the subcortical limbic network nodes, hippocampus and amygdala received 78% of its connections from association networks, including majorly from limbic, default networks and remaining from other networks. Specifically, hippocampus ROIs received connections from perirhinal, entorhinal and parahippocampal cortices^41^, involved in visuospatial processing, encoding spatial maps, episodic and declarative memory processes, face recognition, scene processing, integrating object information with spatiotemporal information. Additionally, convergence occurred from areas involved in higher-order perceptual processing (V2, VMV1, PoI, Area 2, STSda, PGp), processing emotional aspects of auditory, visual, olfactory stimuli and language comprehension (TGd, Areas 47), executive functions (Areas 10, POS, PF). Interestingly, the hippocampus (HIP-Tail) was the only subcortical region that received at least one projection from all cortical RSNs. This is in line with its central role in memory encoding, where the perceptual, emotional, and cognitive dimensions of an experience are bound together within a spatiotemporal framework. Convergence in amygdala occurred from sensory association areas (STGa, STSda, A5, PoI, PH), valuation area integrating internal/visceral states with external information (13l in OFC), (para)limbic areas (TGd, PerC), and language/semantic processing areas (Area 6r, 9m, 44, 47) (**Supplementary Table S3).** Together, this provides evidence for cortical convergence within SLN in relation to limbic functions, including emotion and memory^6^. Overall, the convergence occurred from multiple, widespread, and diverse cortical sites, which appeared segregated along four broad functional-dimensions – sensorimotor integration, motor control, associative, and limbic functions^6,8,9,11^ and mapped onto distinct subcortical networks, resulting into a topographic organization (**Fig. 5D**). The subcortex further supported integration of signals from these segregated circuits via its between-community connections, i.e. connections between nodes of THA, BGN, and SLN.

#### Functional diversity driving cortical regions to converge

To determine how convergence was related to distance and node type, we obtained distributions of distances between the converging cortical node pairs of two types – functionally similar (FS) and functionally diverse (FD). A cortical node pair is FS, if both its nodes belong to a same RSN and is FD, otherwise, i.e. if its nodes belong to distinct RSNs. For this, we computed the Euclidean distance and the proportion convergence for a cortical node pair. Proportion convergence captures how often two nodes project simultaneously to the subcortical nodes with respect to the total convergence. We found that the distributions could be well approximated with normal functions (**Fig. 6**), 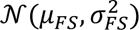 where, 𝜇_𝐹𝑆_ = 27.62 *mm*, 𝜎_𝐹𝑆_ = 11. 1 *mm* for FS node pairs and 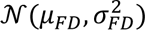 where, 𝜇_𝐹𝐷_ = 8. *mm*, 𝜎_𝐹𝐷_ = 27.70 *mm* for FD node pairs. The proportion convergence strengths of FD nodes were significantly greater than those of FS nodes (paired *t-*test of bin-wise strengths, *t*[16] = 3.84, *p* = 0.0014). Majorly, we found that the cortical regions from functionally diverse communities participated in the converging organization at rest.

**Figure 6.**
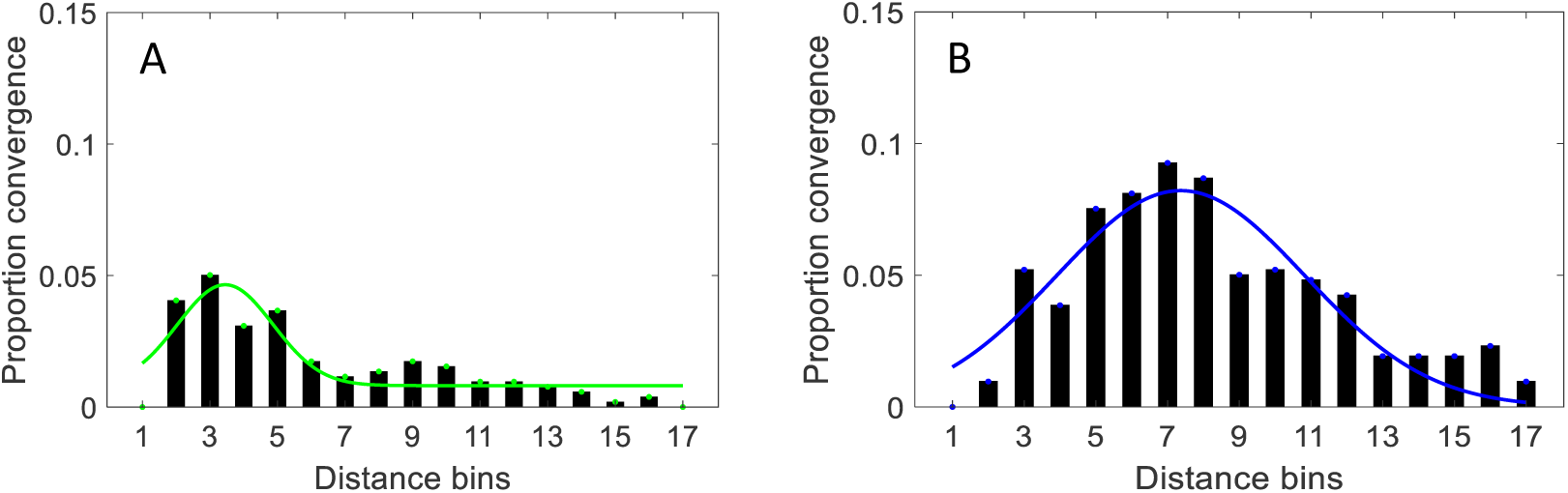
Normal distribution of distances between converging cortical regions. **A** Distribution representing proportion convergence for functionally similar cortical node pairs divided into bins of size 8 mm and approximated using normal function 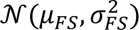 where, 𝜇_𝐹𝑆_ = 27.62 *mm*, 𝜎_𝐹𝑆_ = 11. 1 *mm* (solid green curve). **B** Distribution same as in A for functionally diverse cortical node pairs and approximated using normal function 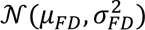 where, 𝜇_𝐹𝐷_ = 8. *mm*, 𝜎_𝐹𝐷_ = 27.70 *mm* (solid blue curve). A cortical node pair is functionally similar, if both its nodes belong to a same resting state network and is functionally diverse, if its nodes belong to distinct resting state networks.

#### Subcortical nodes under targeted attack

We validated the indispensable role of a node in the cortico-subcortical converging organization by assessing the damage caused by an attack on that node, simulated here by removing the node (i.e., all its connections) (**Fig. 7A**). We quantified the potential impact of the attack by measuring its effects on the local information transfer efficiency of the attacked node. The local efficiency (LE) is average efficiency of local neighbourhoods and it shows how efficiently the immediate neighbours of a node communicate when the node is removed^43^. We simulated a targeted attack on 27 subcortical nodes and compared it with random attacks (10,000 simulations) on the converging organization. The neighbourhood of a subcortical node comprises of other subcortical nodes and “converging” cortical nodes. **Fig. 7B** shows the efficiency of individual neighbourhoods due to attack on respective subcortical node. Additionally, we also simulated a more restricted random attack, in which we randomly selected a set of 27 cortical nodes of the converging organization. The latter condition was used to position the role of cortical nodes participating in the converging organization. We found that the local efficiency in case of targeted attack was lesser in comparison to that in any random attack condition (**Fig. 7C**; 𝐿𝐸_𝑡𝑎𝑟𝑔𝑒𝑡_ = 0.58; mean 𝐿𝐸_𝑟𝑎𝑛𝑑𝑜𝑚_ (± SD) = 0.61 (±0.02), *t*[9999] = 132.82, *p* < 0.01; mean 𝐿𝐸_𝑟𝑒𝑠𝑡𝑟𝑎𝑛𝑑_ (± SD) = 0.62 (±0.01), *t*[9999] = 318.37, *p* < 0.01). This indicated a greater and significant impact of subcortex damage than any random damage to the converging organization.

**Figure 7.**
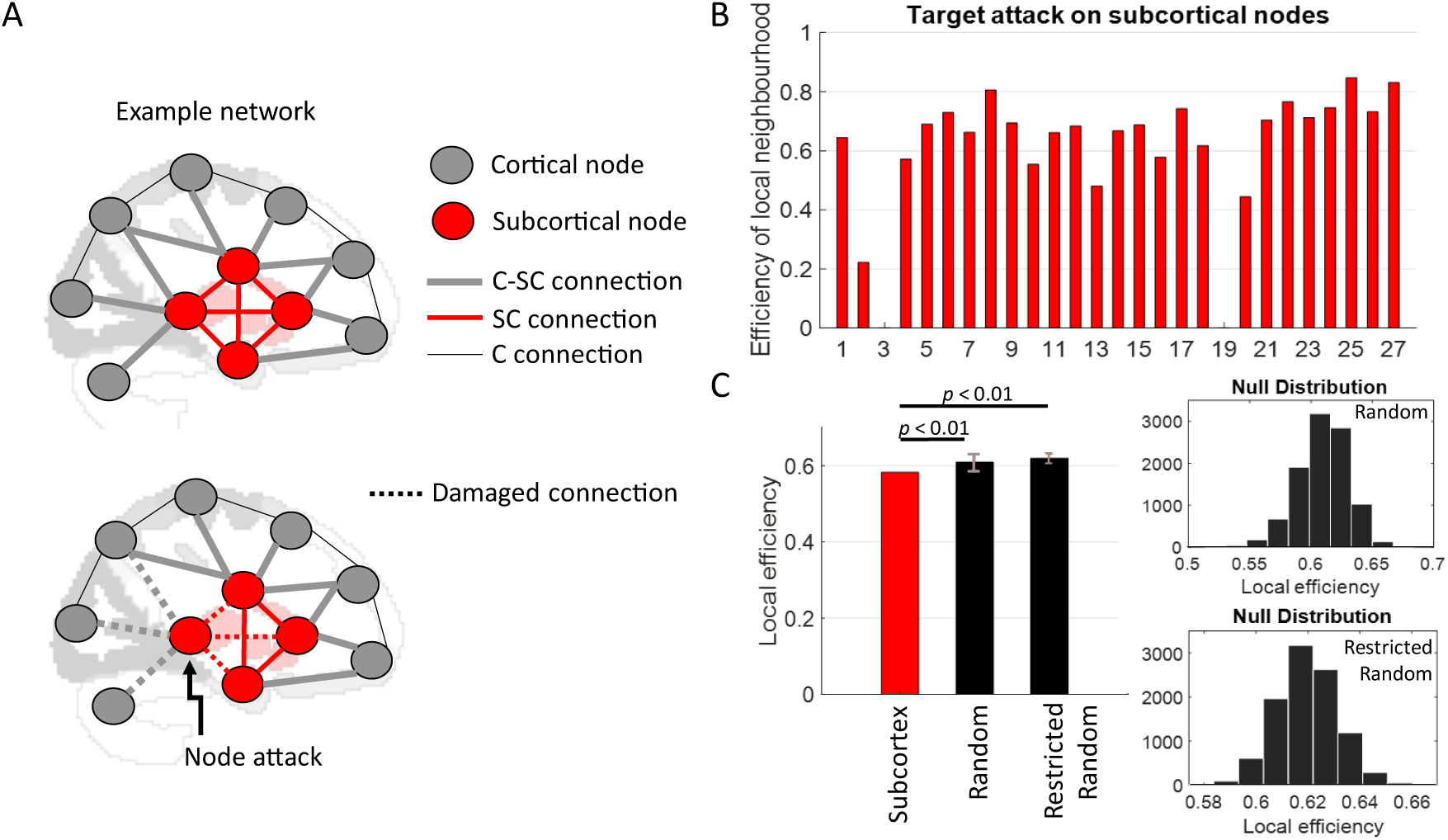
Subcortical nodes under attack. **A** shows an example network with cortical and subcortical nodes as well as different types of connections between – two cortical nodes (C connection), two subcortical nodes (SC connection), and a cortical node and a subcortical node (C-SC) connection. Also shown for illustration is attack on a subcortical node, where all its existing connections are removed (dashed line). **B** Efficiency of neighbourhood of a subcortical node, representing how efficiently the communication occurs between the immediate neighbours of the node, when that node is removed. **C** Local efficiency (LE) in case of targeted and random attack conditions, illustrating greater impact of subcortical damage on the efficiency of cortico-subcortical converging organization in comparison to random attacks (p < 0.01). Also shown are null distributions of LE values from 10000 simulations in case of two random attack conditions. Error bars (in grey) indicate standard deviation of LE values across 10000 random attack simulations.

## Discussion

Distinct cognitive abilities place varying requirements on local segregation and global integration processes in the brain. The resting state organization within the cortex is configured to maintain a dynamic balance between segregation and integration to support diverse “cognitive phenotypes”^14^. The prevalence of resting state research towards cortex, along with technical difficulties intrinsic to subcortical imaging, has caused the subcortex to be an underexplored territory^44^. Availability of high-quality neuroimaging datasets in public-domain^30^ has ushered the exploration of this deep and compact brain structure. To address the gaps in the existing subcortical literature, we answered how resting state subcortex would configure with respect to the functional segregation and integration dynamics. To this end, we determined not only the resting state networks within subcortex but also the cortico-subcortical convergence at rest. We used multi-session resting state fMRI from 92 healthy adults^30^.

### Resting state networks of subcortex

Using partial correlation based functional connectivity and clustering algorithm, we reliably detected three functionally segregated communities within subcortex – thalamic network, basal ganglia network, and subcortical limbic network (**Fig. 2,3**). These RSNs were driven by the well-known specialized functional roles of their nodes. For instance, besides being primarily involved in multisensory/sensorimotor integration^18,45^, thalamic nuclei also involve in cognitive control, working memory, attention^4^. Similarly, the basal ganglia primarily involve in motor control (deciding which actions to allow and which to inhibit) but also contribute to reward processing and executive decision-making in-concert with diverse frontal areas^3,7^. Lastly, amygdala primarily processes emotionally salient stimuli and couples with hippocampus where emotional aspects along with sensory and cognitive dimensions of an experience are bound together for a unified memory encoding^5,6^. Further, the subcortical RSNs reflected not only putative functional boundaries but also anatomical boundaries, which is in line with past studies suggesting that the strong functional connectivity between the nodes of a network is due to direct neuroanatomical connections existing between them^46^.

### Subcortical hubs

Subcortex exhibited equivalent topological characteristics as cortex and contained roughly four times more connectors than non-connectors (**Fig. 4**). We observed that the connectors were positioned radially outward in the periphery of subcortex, including nodes in hippocampus, amygdala, putamen, dorsal caudate nucleus, nucleus accumbens, and dorsal, ventrolateral, posterior thalamus whereas the non-connectors were located inwards in the core of subcortex, including nodes in ventromedial thalamus, ventral caudate nucleus, and globus pallidus. This organization is intuitive given that the connectors participate in multiple communities across whole brain, whereas non-connectors including provincial hubs connect within their community only. Previous work has shown hippocampus, amygdala, putamen, caudate head, and thalamus as connector hubs^4,26,33^. Our results provided new evidence for the existence of “topographical organization” of hubs in subcortex. Overall, this underscores that despite its deep and compact structure, subcortex contributes significantly in global brain communication and in shaping the large-scale dynamics^20^.

### Cortico-subcortical convergence at rest

The main finding of this study is the existence of large-scale cortical convergence within subcortical networks at rest, which overall represented a topographic organization of functionally segregated cortico-subcortical circuits (**Fig. 5**, **Table 1**). These circuits existed along four broad functional dimensions – sensorimotor integration, motor control, associative, and limbic. The sensorimotor circuit majorly mapped onto the thalamic network and provided evidence for existence of communication at rest between brain-wide regions involved in the integration of (multimodal) sensory and motor information^18,45^. This is crucial for higher-order control of global information processing^4^. Further, the associative and motor control circuits mapped onto the basal ganglia network, indicating resting state interaction between regions involved in instrumental learning processes^19,40^. These circuits received sensory inputs via thalamic nodes for refining and limbic inputs via basal ganglia nodes for biasing – the development and control of non-innate actions (sequences) during goal-directed and habitual behaviours^11,40^. Finally, the limbic circuits were mapped onto the subcortical limbic network, indicating resting state communication between regions involved in emotion, memory, and related behaviours^5,47^. Moreover, we found evidence at rest for three distinct but partly overlapping circuits forming the illustrious limbic model^6^ – the parahippocampal-hippocampal circuit associated with spatiotemporal dynamics and memory^5,41^, the default circuit involved in autobiographical memories and introspective thoughts consisting of episodic and semantic memory^48^, and the temporo-amygdala-orbitofrontal circuit involved in integration of sensory signals including visceral sensations and emotion with semantic memory^49^, as emotions are often conveyed and interpreted via language.

Previous resting state neuroimaging studies^24–27^ have adopted a cortico-centric approach and focussed on using cortical network partitions for delineating the network-level activity within subcortex. Our approach is rather novel and subcortico-centric, where we decompose activity with a subcortical region based on region-level activity from both cortical and other subcortical regions. Overall, the cortico-subcortical converging organization at rest demonstrated not only integration between functionally diverse and distant cortical regions but also that subcortex served as an appropriate anatomical substrate for further convergence of information between distinct functional systems (**Fig. 5,6**), due to its key anatomical features – proximity of many smaller nuclei^1^ and progressive compression of pathways^19^. Specifically, the existence of between-community interactions within subcortex represented integration between functionally segregated cortico-subcortical circuits, which were mapped onto distinct subcortical networks. The indispensable role of subcortex for the converging organization was validated by decreased local efficiency within the organization during targeted attack condition in comparison to random attacks (**Fig. 7**). Together, our results reveal how subcortex configures itself at rest in healthy human adults and contributes significantly to orchestrating the whole brain functional segregation and integration dynamics. Future work will focus on revealing any alterations in the resting state configuration of subcortex during distinct task^14^ and diseased^28,29^ states.

## Methods

### Resting state fMRI data

In this study, we examined a publicly available resting state functional neuroimaging dataset from the Human Connectome Project (HCP)^30,50^. We used data from 92 participants (46 males and 46 females, age range = 22 - 35 years, mean (± SD) age = 29.49 (±3.56) years) from the HCP 100 unrelated subjects’ release. All participants provided an informed consent, and the study protocols were approved by the Institutional Review Board at Washington University. Data were collected using a Siemens 3T Connectom Skyra with a 32-channel head coil (detailed acquisition protocols are available here ^50^). Participants were instructed to keep their eyes open and focus on a cross presented on a dark background. Resting state fMRI (rs-fMRI) data were acquired from four scan sessions of approximately 15 minutes each, over a period of two days, using a gradient-echo planar imaging sequence (parameters of imaging: TR =720 ms, TE = 33.1 ms, flip angle = 52°, number of slices = 72, slice thickness = 2mm, 2 mm isotropic voxel resolution, and multiband factor = 8). During each day, oblique axial acquisitions alternated between two phase encoding directions: right-to-left (RL) in one scan session and left-to-right (LR) in the subsequent scan session. We used data from scan sessions with LR phase encoding directions only.

### Preprocessing

We employed the minimally pre-processed, cleaned data (ICA-FIX clean dataset^51,52^. The minimal preprocessing pipeline (MPP) processed fMRI data by removing spatial distortions, realigning scans for subject motion, registering functional scans to the structural data, reducing the bias field, and normalizing the data to a global mean. Further, to maximize spatial correspondence between participants, the timeseries were mapped from a volume space to a standard “gray ordinate” space (more details can be found here^51^; followed by smoothing to regularize the mapping process. The primary goal of MPP was to retain as much relevant information as possible. However, further preprocessing techniques—such as spatial smoothing, temporal filtering, nuisance regression, and scrubbing—can inadvertently remove valuable neural signal activity as highlighted in previous studies^51^. Additionally, there remains ongoing debate regarding the best strategies for these techniques^53,54^, emphasizing the need for careful consideration based on specific research goals and data characteristics. Thus, we refrain from applying any more cleaning procedures than just implemented in ICA-FIX cleaning procedure. The ICA-FIX process removes structured spatial/temporal artefacts and motion fluctuations from the data^52^. All further analyses were carried out using custom-written routines in MATLAB software (Version 9.11.0.1809720 (R2021b) Update 1), including functions from Brain Connectivity Toolbox^55^.

### Whole brain functional parcellations

We used a combination of functional parcellation schemes defining 400 cerebral cortex parcellations (200 per hemisphere) from Schaefer atlas^56^, 26 cerebellar cortex regions from Automated Anatomical Labelling (AAL) atlas (9 bilateral divisions and 8 medial divisions (Vermis))^57^, and 54 subcortex parcellations (27 per hemisphere) from a recent multiscale subcortical atlas^58^. We refer cerebral cortex and cerebellar cortex commonly as cortex, unless otherwise mentioned specifically. **Supplementary Fig. S1** shows 400 cerebral cortex parcellations as part of 7 canonical resting state networks according to^21^, including visual network (VIS), somatomotor network (SOM), dorsal attention network (DAN), ventral attention network (VAN), limbic network (LIM), control network (CON), and default mode network (DMN). Additionally, the figure shows cerebellum (CER) consisting of 26 parcellations and 7 subcortical regions (thalamus, putamen, caudate nucleus, nucleus accumbens, globus pallidus, hippocampus, and amygdala) consisting of 54 parcellations.

### Regions-of-interest

Resting state network organization in human brain is highly symmetrical^21,22,59^. Thus, for further analyses we considered the symmetrical bilateral parcellations for an individual brain area as one region-of-interest (ROI). This resulted into a total of (𝑁 =) 244 ROIs, which included 200 cortical ROIs, 17 cerebellum ROIs, and 27 subcortical ROIs. For each participant, we extracted the summary (mean) time courses for each ROI from the ICA-FIX cleaned data. We discarded the first 100 volumes (approx. 72s) from each of the two LR phase encoding scan sessions (completed over two days) because the first few volumes of a functional acquisition contain large signal changes which stabilise as the tissues reach steady state^60^. Further, the time courses from individual sessions were mean-centred to remove the mean intensity difference across sessions and then concatenated^25^.

### Functional connectivity

To reveal underlying network organization at rest and compute network-based metrics, a whole brain functional connectivity^61^ (FC) matrix was determined based on a common statistical modelling approach^31,62,63^. We quantified the *partial* pair-wise correlation between ROIs by using a multiple linear regression model. Partial correlation captures unique dependence between any two nodes while controlling for the effects of all other nodes in consideration (i.e. subcortical as well as cortical nodes). For each participant, we estimated the mean temporal activity from a ROI, say *X*_*i*_, based on the activity of remaining 𝑁 − 1 (= 243) ROIs. This was done by including the remaining ‘*N* − 1’ time courses as regressors in the model. In this way, activity in a region *X*_*j*_ explained the activity or captured the partial variance in the estimated time course *X*_*i*_ uniquely due to itself and the shared variance due to remaining regressors, say *X*_*k*_, was not attributed to the regressor *X*_*j*_, where, *i*, *j*, *k* ∈ {1,2, …, *N*} and *i* ≠ *j* ≠ *k*. Thus, the parameter estimates 𝛽_*j*,*i*_ (coefficient of regressor *X*_*j*_ in the model estimating *X*_*i*_) provided a weighted measure of unique or partial influence of the activity in ROI *X*_*j*_ on the activity in ROI *X*_*i*_, by controlling for the influence from other ROIs *X*_*k*_. The higher the magnitude of the parameter estimates, the greater the correlation between the regions. Using the above approach helped controlling for any false positives. Self-dependence was not modelled based on the method used in^31,62^. No intercept term was included in the regression model as the regional time courses were mean-centred. For each participant, we repeated the above estimation procedure for each of the *N* whole brain ROIs.

*For group-level significance*, individual parameter estimates from each participant were subjected to one-sample *t*-test (a random effects analysis method^64^). For all reported results, we employed a significance threshold of *p <* 0.01, Bonferroni corrected for multiple comparisons (equal to number of regressors in the model as the activities of the nodes are not fully independent^32,65^). Finally, we computed a group-level whole brain *binary* functional connectivity matrix 𝐶 (of size *N* × *N*), where the element 𝐶_*j*,*i*_ is 1, if activity in ROI *X*_*j*_ had significant influence on the activity in ROI *X*_*i*_; otherwise, 0. The connectivity matrix was asymmetrical as 𝐶_*j*,*i*_ may or may not equal 𝐶_*i*,*j*_. We also computed a group-level whole brain *weighted* functional connectivity matrix 𝑤𝐶, where the elements of the matrix were significant parameter estimates from the models, i.e., 𝑤𝐶_*j*,*i*_ is 𝛽_*j*,*i*_, if activity in ROI *X*_*j*_ had significant influence on the activity in ROI *X*_*i*_; otherwise, 0. Thereafter, we normalized the weights in the matrix, such that they ranged between −1 𝑡𝑜 1.

### Subcortical community detection

#### Louvain clustering algorithm

To determine whether and how parcellations in subcortex formed functionally segregated communities at rest, we employed the Louvain clustering algorithm^66^. A community consists of a subset of unique nodes, which have stronger connections within the community in comparison to outside than expected in a random “null” network, resulting into nonoverlapping partitions or modules of a network, for more details refer^67^. The degree of modularity in a network can be characterized by the *Q* index^68,69^. The Louvain algorithm operates as follows: it begins by identifying smaller communities through optimization of a modularity function *Q*, which is primarily based on the connectivity matrix of the network nodes, a resolution parameter 𝛾, and an initial random community assignment. Next, it merges these smaller communities into larger ones, creating a new network. This iterative process continues until there are only minimal changes in the value of modularity index *Q*. The algorithm aims on enhancing the strength of connections within a community rather than the connections between the communities^66^.

We determined the functional networks in subcortex based on the group-level binary functional connectivity matrix of 27 subcortical ROIs (of size 27 x 27) computed in previous section. The resolution parameter 𝛾 was set to standard value of 1 and 1000 random initializations of community assignment were made. The resolution parameter 𝛾 controls the number of community partitions, where 𝛾 < 1 partitions a network into fewer (but larger) nonoverlapping communities and 𝛾 > 1 into more (but smaller) nonoverlapping communities than standard (𝛾 = 1). Further, even for a set 𝛾 value there are multiple possible solutions that optimize the modularity function *Q*, resulting into partitioning outcomes/solutions that vary across different random initializations of community assignment. To control for this variability, we performed clustering 1000 times each starting with a random community assignment and thereafter, employed a consensus approach across all 1000 partitioning outcomes. The consensus approach computed one final partitioning scheme using the Louvain algorithm on an agreement matrix (27 x 27; like a connectivity matrix) and a resolution parameter 𝜏. The agreement matrix contained the probability of each pair of nodes to be assigned to the same community across all 1000 outcomes. The consensus approach used a parameter 𝜏 which served as a threshold value for the agreement matrix to control the resolution of final partitioning scheme. We set the value of 𝜏 to unity, so that the pair of nodes which have been consistently partitioned (with probability 1.0) into the same community across all 1000 outcomes were included into the computation of final partitioning scheme. We refer to it as the “representative” resting state network organization in subcortex.

#### Silhouette score

The silhouette score 𝑆_*i*_ for a node *i* evaluates its closeness (a measure of distance in terms of connectivity strength) to the other nodes in the same community compared to the nodes in the next nearest community^70^. The higher the connectivity strength between two nodes, the closer the two nodes. The score was computed using weighted partial connectivity matrix *wC* of subcortical nodes (27 x 27), as shown below:

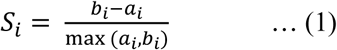

where, 𝑎_*i*_ is the average distance from node *i* to the other nodes in the same community as node *i*, and 𝑏_*i*_ is the minimum average distance from node *i* to the nodes in a different community, minimized over subcortical communities. The silhouette score for each node ranges from –1 to 1, where a positive value signifies greater confidence in the current community assignment. Conversely, a negative value indicates stronger association on average to the next nearest community than its assigned community. If most nodes have a high silhouette score, then the clustering solution is appropriate. If many nodes have a low or negative score, then the clustering solution indicates too many or too few communities. Next, we performed a more rigorous validation of our results.

#### Statistical validation

Statistics-based network analysis^32,71^ was used to validate whether the characteristics of the segregated organization in subcortex were significantly different than chance. For this, a comparison of real network was done with 1000 random “null” networks. The random networks were generated by rewiring connections between nodes while maintaining the same degree distribution as our “representative” subcortical network organization at rest (obtained above). This is a commonly used approach for network validation^15,32,33,72^. Characteristic graph theory metrics were computed for the real and random networks both at node-level and network-level, including measures of segregation (within-module degree (WMD), modularity index 𝑄) and those of integration (participation coefficient (PC), characteristic path length (CPL)). We included integration metrics to check whether our community assignment results were affected by nodes with substantial between-module connections that contradict the hypothesis of segregated RSNs at rest.

For computation of all graph metrics used here, we considered the group-level binary partial connectivity matrix (27 x 27) as a binary directed graph. We briefly describe each metric next (for implementations and more details refer^55,73^. The measures of segregation characterize the community structure and communication within a community, where within-module degree *wmd*_*i*_ is the number of connections *k*_*i*𝑚_ of node *i* in its own community 𝑚. A corresponding z-score metric 𝑧_*i*_ shows how well-connected is node *i* within its own community 𝑚 in comparison to other nodes of its own community, as shown below:

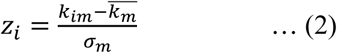

where, *k*_*i*𝑚_ denotes the number of within-module connections (bidirectional – incoming and outgoing) of a node *i* in its community 𝑚, *k*_𝑚_ and 𝜎_𝑚_ are the average and standard deviation values across the number of within-module connections of all nodes in community 𝑚, respectively. The modularity index 𝑄, a network-level segregation metric, indicates on average how well-connected are the communities within themselves in comparison to each other ^67^. A 𝑄 value ranges from 0 to 1, where a value close to 0 suggests that the network resembles a random network, while a value close to 1 indicates a strong community structure.

The measures of integration characterize communication between the communities, where participation coefficient 𝑃_*i*_ indicates how well-connected is a node to all communities of the network, as shown below:

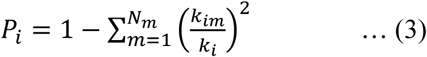

where, *N*_𝑚_ is the number of communities, please note, in the subcortex only, *k*_*i*_ is the degree of node *i* (total number of bidirectional connections – incoming and outgoing) and *k*_*i*𝑚_ is the number of connections between node *i* and nodes of any community 𝑚. If a node’s connections are entirely restricted to its own community, its PC is 0; while a node with high PC (closer to 1) connects to multiple communities. The characteristic path length 𝜆, a network-level integration metric, is the average shortest distance between all pairs of nodes in a connected network. A lower CPL value indicates higher integration between communities resembling a random network.

For each of the four characteristic graph metrics (i.e., within-module degree, modularity index, participation coefficient, and characteristic path length), we obtained a “null” distribution of metric coefficients from a population of 1000 random networks. Please note that in case of a node-level metric, say WMD, 27 null distributions were computed each corresponding to the 27 nodes in the subcortex. Then, using one-sample *t*-tests we checked whether the real network’s metric values were significantly different from their respective null distributions. For all reported results, we employed a significance threshold of *p <* 0.01, Bonferroni corrected for multiple comparisons as the activities of the nodes are not fully independent^32,65^.

#### Topology of subcortex

To understand the role of subcortex in local segregation and global integration dynamics, we examined its topology using network science metrics^73^. Specifically, we identified four distinct types of nodes in subcortex, including connector hubs, non-hub connectors, provincial hubs and non-hubs. Connectors hubs are brain regions which have many connections both within their community and outside with multiple communities and non-hub connectors are regions with more connections in other communities in comparison to connections within their community, whereas provincial hubs are regions which have many connections within their own community^15,33,72^. We classified the 27 subcortical nodes based on their participation coefficient and WMD z-score (see equations (2), (3) in previous section). We used a total of (*N*_𝑚_ =) 11 whole brain community partitions (covering *N =* 244 ROIs), including the 3 subcortical and 8 cortical communities, and group-level binary FC matrix (of size 244 x 244). A node *i* was classified as “connector hub” if its participation coefficient 𝑃_*i*_ > 0. and WMD z-score 𝑧_*i*_ > 0; as “non-hub connector” if 𝑃_*i*_ > 0. and 𝑧_*i*_ ≤ 0; as “provincial hub” if 𝑃_*i*_ ≤ 0. and 𝑧_*i*_ > 0; and as “non-hub” if 𝑃_*i*_ ≤ 0. and 𝑧_*i*_ ≤ 0 (using equations (2), (3) with *N*_𝑚_ = 11)^34,73^. The percentage of a particular node type (say, connectors in subcortex) was calculated by dividing the number of that node type with the total number of nodes (in the subcortex).

Further, we compared the topology of subcortex with that of cortex based on the following characteristic graph metrics^73^ – degree (absolute number of connections), within-module degree, connection density (degree per voxel), clustering coefficient (of a node is local graph metric indicates how well the neighbours of that node are connected), and participation coefficient (a global graph metric). In addition to comparing the absolute number of connections (degree) across node types in subcortex and cortex, we also compared their connection densities to test whether subcortex can support functional integration equivalent to cortex, despite its smaller nuclei structures. The connection density of a node was measured by dividing its degree with the size of the region, i.e. the number of isotropic voxels (2mm x 2mm x 2mm) contained in the region. Some of the metrics are described in previous section.

### Cortico-subcortical converging organization

#### Convergence and distance metrics

To determine the extent of convergence in the subcortex, we computed the absolute number *N*^𝑐𝑜𝑛𝑣^ of cortical ROIs converging (via bidirectional connections) onto a subcortical ROI within the subcortex as below:

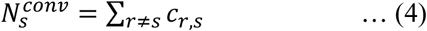

Where, 𝑐_𝑟,𝑠_ = 1, if a bidirectional connectivity exists between cortical node 𝑟 and subcortical node 𝑠 and 𝑐_𝑟,𝑠_ = 0 otherwise, computed using the group-level whole brain binary directed graph 𝐶 = {𝐶_𝑟,𝑠_} (of size 244 x 244) at a significance threshold of *p <* 0.01, Bonferroni corrected for multiple comparisons. We considered bi-directional connectivity between nodes because “dual dyad” motif (a subgraph of three nodes having two sets of bidirectional connections joining at a third single node) have been found to be present in abundance in larger network as compared to other possible subgraphs^74^. Motifs are building blocks of a larger network and unfold underlying statistical properties of structural and functional brain organization^74,75^. Also, it seems biologically plausible to have a dual dyad motif in abundance, given that not only integration but also distribution of information is crucial for efficient brain function.

To determine how convergence between the cortical nodes was related to the distance between the nodes and the types of nodes (functionally similar or diverse) in cortex, we computed proportion convergence 𝑂_*i*,*j*_ between cortical nodes *i* and *j*. In past, a similar analysis just based on distance has been done for tract-tracing data from non-human primates^17^ but is still missing for non-invasive human resting state data. We categorized a cortical node pair as functionally similar, if both its nodes belong to a same RSN and as functionally diverse, if its nodes belong to distinct RSNs. The metric 𝑂_*i*,*j*_ is the proportion of convergence for the cortical node pair (*i*, *j*) with respect to the total convergence, calculated as below^17^:

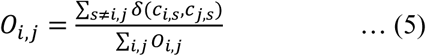

Where, 𝛿(𝑥, 𝑦) is the Kronecker delta that is 1 if x is equal to y and 0 otherwise, 𝑠 denotes a subcortical node, 𝑐_𝑟,𝑠_ = 1, if a bidirectional connectivity exists between cortical node 𝑟 and subcortical node 𝑠, and 0 otherwise, as aforementioned. The metric value lies between 0 to 1 and captures the frequency of convergence, i.e., a higher metric value indicates that a cortical node pair converges at multiple subcortical nodes.

Next, we computed the Euclidean distance (in mm) between the cortical parcellations *i* and *j* as below:

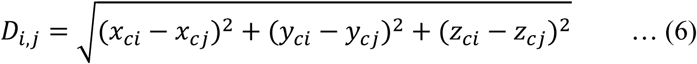

Where, (𝑥_𝑐𝑟_, 𝑦_𝑐𝑟_, 𝑧_𝑐𝑟_) denotes the coordinates of the centroid of a brain parcellation 𝑟. Since our ROIs (nodes) are formed using bilateral parcellations, to compute distance between two cortical ROIs we averaged over the distance between respective parcellations in left hemisphere and the distance in right hemisphere. To obtain distribution of distances, we divided the cortical node pairs into bins of equal distance range, i.e. 8 mm bin sizes, and summed the proportion convergence of the cortical node pairs falling into a particular bin.

#### Node attack and local efficiency

To quantify the potential effects of attack on a set of nodes, we measured the local efficiency of the attacked nodes^43^. The local efficiency (LE) is average efficiency of *N*_𝐺_ local subgraphs or neighbourhoods. The efficiency 𝐸(𝐺_*i*_) of a local neighbourhood 𝐺_*i*_ is the strength of communication among the immediate neighbours of node *i*, when node *i* is removed, computed as below:

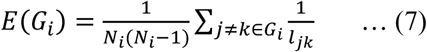

Where, *i* = 1, …, *N*_𝐺_, 𝐺_*i*_ is the neighbourhood of node *i*, *N*_*i*_ is the number of nodes in neighbourhood 𝐺_*i*_, 𝑙_*jk*_ is the shortest path length between nodes *j*, *k* (≠ *i*) of the neighbourhood 𝐺_*i*_. A path in a network graph is defined as the minimum number of distinct edges that must be traversed to pass from one node to another in the network^32,73^. It is a measure of how efficiently two nodes can transmit information to each other. A lesser LE value indicates, on average, poor communication in the local neighbourhoods and thus, a greater impact of attack.

We simulated two types of attacks^76^: (1) a targeted attack which refers to damage inflicted on a particular set of nodes/connections in the network, and (2) a random attack which refers to damage inflicted on a set of randomly selected nodes/connections in the network. Further to compare the effects of targeted attack, we generated a null distribution of local efficiency values by simulating 10000 random attacks. Specifically, we used a one-sample t-test with null hypothesis that data of 10000 LE values comes from a distribution with mean equal to the local efficiency of target attack. A rejection of null hypothesis at p < 0.01 would indicate that there is significant difference in the effects of targeted attack and that of random attack.

## Authors’ contributions

DJ – Conceptualization, data curation, clustering analysis, writing first draft, reviewing & editing manuscript. SD – Conceptualization, convergence analysis, writing first draft, reviewing & editing manuscript.

## Data and code availability

The dataset used in the current study is publicly available from Human Connectome Project. The codes used in the current study will be deposited in Open Science Framework after acceptance of the manuscript.

