## Supplementary Information for "Cortico-subcortical converging organization at rest"

#### Whole brain functional parcellations

We used a combination of functional parcellation schemes defining 400 cerebral cortex parcellations (200 per hemisphere) from Schaefer atlas[1], 26 cerebellar cortex regions from Automated Anatomical Labelling (AAL) atlas (9 bilateral divisions and 8 medial divisions (Vermis))[2], and 54 subcortex parcellations (27 per hemisphere) from a recent multiscale subcortical atlas[3], shown in **Fig. S1**.

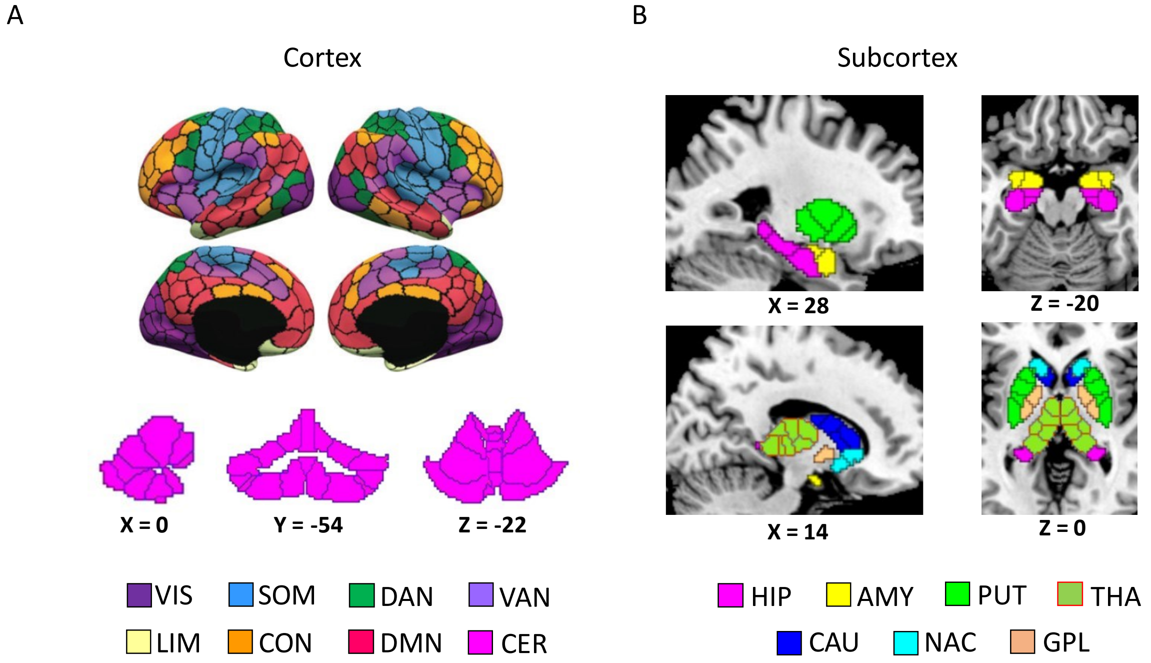


**Figure S1. Whole brain functional parcellations. A** 200 functional parcellations per hemisphere of cerebral cortex shown here as part of existing 7 canonical resting state networks[4]. The RSNs include visual network (VIS), somatomotor network (SOM), dorsal attention network (DAN), ventral attention network (VAN), limbic network (LIM), control network (CON), and default mode network (DMN). Additionally, shown are 26 parcellations of cerebellum (CER). **B** 27 functional parcellations per hemisphere of subcortex shown here as part of 7 subcortical regions, including thalamus (THA), putamen (PUT), caudate nucleus (CAU), nucleus accumbens (NAC), globus pallidus (GPL), hippocampus (HIP), and amygdala (AMY).

### Group-level functional connectivity based on full correlation

For each participant, we first computed the Pearson's correlation coefficient between the pre-processed time series of the 27 subcortical ROIs. This resulted into a 27 x 27 correlation matrix. To obtain the group-level matrix, we did the following based on[5]: We Z-transformed the individual correlation matrices, averaged across participants, and back-transformed into Pearson correlation coefficients $\boldsymbol{r}$. This resulted into group-level functional connectivity based on pair-wise full correlations between the subcortical ROIs (see **Fig. S2**). Again, we do not show here the self-correlations which are of unit strength. We found that the block-like patterns were missing (as in **Fig. 2** of main text), which indicated that the full correlation values did not reflect the true or unique dependence between the ROIs. The shared variance due to other whole brain ROIs was reflected in the form of higher correlation between any two subcortical ROIs, including even between those ROIS which belonged to distinct RSNs ($\boldsymbol{r} > 0$, $t[(\frac{Ns(Ns-1)}{2}) - 1] = 20.19$, $p < 0.01$, Bonferroni corrected, where $Ns = 27$). Thus, this indicated that partial correlation method detected true connectivity between the regions, and in turn the true modular organization in subcortex.

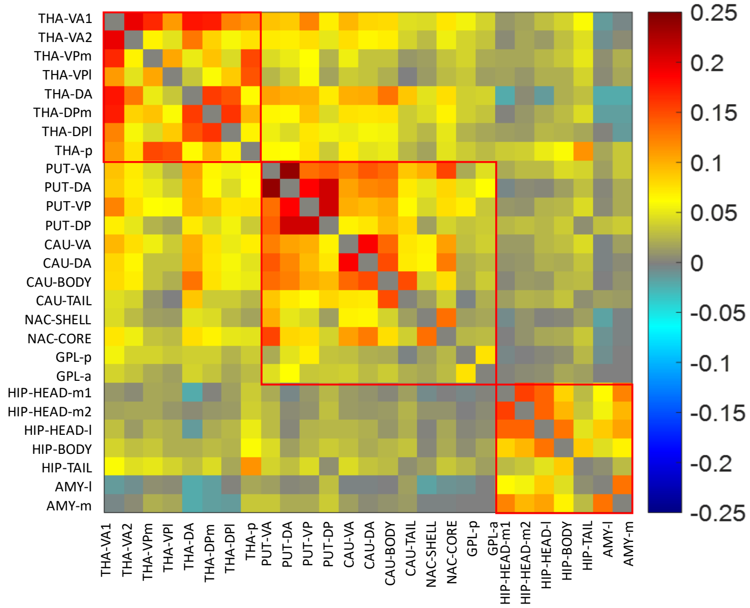


**Figure S2. Group-level ROI-by-ROI functional connectivity matrix based on full correlation.** For comparison, we sort the matrix here according to the three communities identified previously within the subcortex using partial correlation, i.e. the thalamic network (THA), the basal ganglia network (BGN), and the subcortical limbic network (SLN), (ROIs: V – ventral, D – dorsal, A or a – anterior, P or p – posterior, l – lateral, m – medial). The full correlation values range between -0.25 to 0.25.

### Statistical validation results of subcortical community organization

**Table S1** shows one-sample *t*-test statistics for a comparison of within-module degree (WMD) and participation coefficient (PC) across subcortical and random nodes. We observed a significant difference between the two topologies (subcortex and random), where the subcortical nodes were well-connected to nodes of their own community in comparison to nodes of other subcortical communities. This was indicated by higher WMD values and lower PC values of subcortical nodes than their counterparts in random network (null distribution generated using 1000 simulations).

**Table S1. Node-wise statistics for characteristic graph metrics**

| Subcortex ROIs | T(999)* Within-module degree | T(999)* Participation coefficient |
|---|---|---|
| **THA-VA1** | 139.25 | -176.82 |
| **THA-VA2** | 70.71 | -47.04 |
| **THA-VPm** | 115.11 | -117.66 |
| **THA-VPl** | 109.99 | -106.42 |
| **THA-DA** | -5.420 | -55.49 |
| **THA-DPm** | 127.75 | -141.05 |
| **THA-DPl** | 128.97 | -144.6 |
| **THA-p** | 76.17 | -57.33 |
| **PUT-VA** | 174.1 | -270.86 |
| **PUT-DA** | 124.07 | -149.61 |
| **PUT-VP** | 112.34 | -123.49 |
| **PUT-DP** | 131.32 | -155.54 |
| **CAU-VA** | 141.94 | -190.89 |
| **CAU-DA** | 144.46 | -183.62 |
| **CAU-Body** | 137.21 | -172.41 |
| **CAU-Tail** | 129.11 | -158.68 |
| **NAC-Shell** | 81.14 | -65.61 |
| **NAC-Core** | 125.7 | -130.95 |
| **GPL-p** | 94.39 | -75.72 |
| **GPL-a** | 71.40 | -45.04 |
| **HIP-Head-m1** | 96.37 | -73.67 |
| **HIP-Head-m2** | 94.73 | -91.68 |

| | | |
|---|---|---|
| **HIP-Head-l** | 140.29 | -172.92 |
| **HIP-Body** | 77.90 | -61.22 |
| **HIP-Tail** | 56.87 | -33.44 |
| **AMY-l** | 70.54 | -44.33 |
| **AMY-m** | 58.63 | -67.28 |

*corr. P-value < 0.01

### Comparison of topology of subcortex and cortex

We compared the topology of subcortex with that of cortex based on characteristic graph metrics (see **Table S2** below). We found that the nodes in subcortex had significantly lower (within-module) degree than the nodes in cortex ($p < 0.01$ in a two-sample $t$-test). One of the reasons for this could be smaller nuclei structures in subcortex[6] receiving fewer number of connections in comparison to cortical regions with larger sizes. Thus, for a fair comparison we computed connection density metric for each node which was measured by dividing the degree of a node with its volume (i.e., the number of voxels contained in the ROI). Strikingly, we found the connection density values across subcortical ROIs to be greater than cortical ROIs; however, the difference did not reach significance ($p = 0.26$). Similarly, we found no difference between the subcortex and cortex for both clustering coefficients ($p = 0.02$) and participation coefficients ($p = 0.90$). Thus, our results here show that despite of its smaller nuclei structures, subcortex supports both functional segregation and integration equivalent to the cortex.

**Table S2. Graph metrics of nodes in subcortex and cortex**

| **Graph metrics** [a] | **Subcortex (27 nodes)** | **Cortex (217 nodes)** | **T(242)** |
|---|---|---|---|
| **Degree** [b] | 24.89 (±11.05) | 54 (±21.45) | -6.93 (p < 0.01) |
| **Within-module degree** [b] | 10.59 (±3.85) | 23.78 (±11.69) | -5.81 (p < 0.01) |
| **Connection density** | 0.20 (±0.15) | 0.18 (±0.08) | 1.14 (p = 0.26) |
| **Clustering coefficient** | 0.38 (±0.15) | 0.32 (±0.11) | 2.45 (p = 0.02) |
| **Participation coefficient** [c] | 0.61 (±0.22) | 0.62 (±0.19) | -0.12 (p = 0.90) |

[a] Values represent mean (±SD) across nodes, [b] Significant difference between graph metrics of nodes in subcortex and cortex at $p < 0.01$ in a two-sample t-test with equal means and equal but unknown variances, [c] Please note that the PCs have been calculated with respect to whole brain communities.

### Functional roles of converging cortical ROIs

**Table S3** presents in detail the functional roles of cortical ROIs converging onto each of the 27 subcortical ROIs.

**Table S3. Functional roles of converging cortical ROIs**

| ROI index | Subcortical ROI | Converging cortical ROIs* | Function |
|---|---|---|---|
| **1** | THA-VA1 | V1 | Area V1 processes basic features of visual inputs. V1 cells are orientation and direction selective, respond to binocularity and binocular disparity, involve in initial detection of motion. V1 contains retinotopic maps of the visual field[7]. |
| | | Area 4 (M1) | Area 4 controls voluntary movements of specific body parts, responsible for reaching movements of the upper limb, controls breathing, blinking, and learned motor sequences[8]. |
| | | Area p24 (PreACC) | Area p24 is involved in interoception, inhibition of action[9]. |
| | | CER | The cerebellum regulates motor movements and controls balance. It helps coordinate gait, maintain posture, regulate muscle tone, and manage voluntary muscle activity, although it cannot initiate muscle contractions. The vermis cortex oversees the movements of the trunk, including the neck, shoulders, thorax, abdomen, and hips. The intermediate zone of the cerebellar hemispheres, located next to the vermis, controls the muscles of the distal extremities. The outermost part of each cerebellar hemisphere is involved in planning sequential body movements and in the conscious evaluation of movement errors[10]. |
| **2** | THA-VA2 | - | |
| **3** | THA-VPm | - | |
| **4** | THA-VPl | OP1 (S2) | OP1 is part of secondary sensory cortex located on parietal operculum. It contributes to somatosensory processing tasks such as pain |

| | | | |
|---|---|---|---|
| | | | recognition, tactile attention, and working memory[11,12]. |
| | | Area 3b (S1) | Area 3b processes tactile information, such as pressure, vibration, and texture, from the body's skin acting as the initial site of activation during tactile stimulation[13]. It contributes to fine tactile discrimination by localizing sensations on the skin from specific parts of the body (such as the hands, lips, and face) and distinguishing their specific characteristics, in contrast to areas 1 and 2 which are responsible for processing more generalized receptive fields[14]. Area 3b also processes nociceptive stimuli[15] and overall is crucial for perceiving and interpreting tactile sensations, enabling the brain to recognize and respond to touch stimuli from the body[16]. |
| | | Area 2 (S1) | Area 2 is involved in the higher-order processing of sensory information. It integrates sensory input related to touch with proprioception (the sense of body position) and helps in creating a more complex understanding of the body's position and movement in space. Area 2 processes information not only from the skin but also from muscles and joints, allowing it to contribute to the perception of object size, shape, texture, and movement. It helps in constructing a more abstract representation of the body in space[16]. |
| **5** | THA-DA | Area 8BM (dmPFC) | Area 8BM facilitates overall motor planning, including oculomotor control, speech motor programming and executes goal-directed behaviors[17]. |
| | | CER | As above |
| **6** | THA-DPm | Area 3b (S1) | As above |
| | | Area 8BM (dmPFC) | As above |
| | | CER | As above |
| **7** | THA-DPl | V1 | As above |
| | | A1 | Area A1 processes the basic features of sound, such as frequency, pitch, volume for sound |

| | | | |
|---|---|---|---|
| | | | perception and also contains tonotopic maps describing distribution of frequency preference[18]. |
| | | CER | As above |
| **8** | THA-p | V2 | Area V2 integrates information from V1, leading to more complex response patterns to objects. V2 cells respond to variations in color, spatial frequency, moderately complex patterns, object orientation, and the distinction of stimulus from background[19]. |
| | | POS1 (VC) | POS1 area involves in mental navigation, scene perception, and working memory related to place images[20,21]. It shows greater activation when processing socially interactive objects compared to randomly moving geometric shapes[22]. |
| | | Area 2 (S1) | As above |
| | | PF (IPL) | Inferior parietal lobule comprises of 7 cytoarchitectonically defined areas PFt, PFop, PF, PFm, PFcm, PGa, and PGp[23]. Anterior human IPL areas, including PFt, Pfop, PFcm, involve primarily in somatosensory processing; and PF in processing visuo-motor and visual object information, multimodal body image representations[24], also in motor planning and action-related functions and is part of the human mirror neuron system[25,26]. PF involves in action observation and imitation, for instance handling of tools for moving object[26,27]. |
| | | CER | As above |
| **9** | PUT-VA | FOP | The frontal operculum controls the movements required for speech articulation, including the planning and coordination of movements for precise pronunciation and language comprehension, i.e., helps with understanding the syntactical and grammatical aspects of language[28,29]. |
| | | Area a24pr (aMCC) | Area a24pr is part of anterior MCC, which coordinates fear and avoidance behavior via skeletomotor regulation and motor response selection during painful stimulation[30,31]. |

| | | | |
|---|---|---|---|
| | | Area a47r (vlPFC) | Area 47 involves in top-down semantic retrieval and executive control in language processing[32,33]. In addition to its connection with Broca's area, Area 47 is also considered part of the broader "Broca's complex," which also includes Brodmann areas 44, 46, 47, and the mesial supplementary motor area of 6. Together, these regions form a frontal-subcortical circuit[34]. |
| | | Area 9m (dmPFC) | Area 9m is a subdivision of area 9, which involves in cognitive control, executive functions, working memory, cognitive flexibility, and response inhibition[35] |
| | | Area p24 (PreACC) | As above |
| **10** | PUT-DA | Area 4 (M1) | As above |
| | | SCEF (PMC) | SCEF is higher-order oculomotor control area, which involves in evaluating all potential oculomotor actions to guide primary oculomotor sites during goal-directed behavior[36]. |
| **11** | PUT-VP | Area 55b (PMC) | Area 55b is a premotor area and associates with both speech production[37] and music rhythm attunement[38]. |
| | | Area 4 (M1) | As above |
| | | CER | As above |
| **12** | PUT-DP | Area 4 (M1) | As above |
| | | PoI2 | Posterior insular cortex involves[39] in multimodal sensory processing and somatosensory control[40] thermosensory, chemosensory, and pain perception[41], and speech[42]. |
| | | FOP | As above |
| | | Area 6ma (SMA) | The supplementary motor area has four broad subdivisions – SFL, SCEF, 6ma, and 6mp. SMA involves[43,44] in motor planning, initiation and coordination of internally-motivated movements, including limb movement and speech production and externally-cued movements, including reaching, grasping, action selection and inhibition. Specifically, 6ma shows greater |

| | | | |
|---|---|---|---|
| | | | activity compared to SFL, 6mp during visual instruction cues[45]. |
| | | CER | As above |
| **13** | CAU-VA | FOP | As above |
| **14** | CAU-DA | SCEF (SMA) | As above |
| | | Area a10p (dlPFC) | Area a10p involves in episodic memory, abstract cognitive function, and relates to increasing difficulty levels during working memory tasks[46,47]. |
| | | Area p24 (PreACC) | As above |
| | | SFL (SMA) | The supplementary motor area has four broad subdivisions – SFL, SCEF, 6ma, and 6mp. SMA involves[43,44] in motor planning, initiation and coordination of internally-motivated movements, including limb movement and speech production and externally-cued movements, including reaching, grasping, action selection and inhibition. Specifically, SFL shows greater activity during story-listening, object-matching, and social-interaction scenarios[45]. |
| | | CER | As above |
| **15** | CAU-Body | AVI | Anterior insular cortex involves[39] in visceral sensory processing, autonomic control, and interoception[48], cognitive control and emotion processing[49], empathy and social cognition, risky decision-making[50], attention and saliency processing[51]. |
| | | Area 8BM (dmPFC) | As above |
| | | Area 44 (IFG) | Area 44 converts abstract representations from prefrontal cortex into detailed representations that assist in guiding verbal and manual actions[52]. In addition to its connection with Broca's area, Area 44 is also considered part of the broader "Broca's complex," which also includes Brodmann areas 45, 46, 47, and the mesial supplementary motor area of 6. Together, these regions form a frontal-subcortical circuit[34]. |

| | | | |
|---|---|---|---|
| | | CER | As above |
| **16** | CAU-Tail | STGa | STGa is an auditory association area and involves in perceptual and conceptual acoustic sounds processing[53], language processing and socioemotional processing[54,55]. |
| | | CER | As above |
| **17** | NAC-Shell | AVI | As above |
| | | Area 9m (dmPFC) | As above |
| | | Area p24 (PreACC) | As above |
| | | CER | As above |
| **18** | NAC-Core | Area 9/46d (dlPFC) | Area 9-46d, involves in cognitive control processes, such as the monitoring or tagging of recent information rather than passive maintenance of information[35]. |
| | | Area 9m (dmPFC) | As above |
| | | Area p24 (PreACC) | As above |
| **19** | GPL-p | - | |
| **20** | GPL-a | CER | As above |
| **21** | HIP-Head-m1 | PHC | The parahippocampal cortex processes "visuospatial and contextual information in the service of episodic memory or memory for the events of our daily lives". It shows greater activation during place/scene recognition rather than face recognition[56]. |
| | | POS1 (VC) | POS1 area involves in mental navigation, scene perception, and working memory related to place images[20,21]. It shows greater activation when processing socially interactive objects compared to randomly moving geometric shapes[22]. |
| | | PerC | The perirhinal cortex involves in integrating semantic information to support object identification, integrating object information |

| | | | |
|---|---|---|---|
| | | | with spatiotemporal information and further transferring this knowledge to declarative memories in hippocampus via entorhinal cortex[57]. It shows greater activation during selective face recognition and also known as "anterior temporal face patch"[58,59]. |
| | | EntC | The entorhinal cortex involves in "both the rapid encoding of new associations and in the consolidation of memory in connection with the medial prefrontal cortex"[60]. |
| | | STSda | STS involves diverse functions, including speech, motion and facial processing. Additionally, it shows activation in response to unimodal visual and audio inputs and also performs audiovisual integration. Along with other higher-order auditory areas A4, A5, and STGa, it involves in semantic processing of audio information and thus, considered as auditory association cortex[61]. |
| | | PGp (IPL) | Inferior parietal lobule comprises of 7 cytoarchitectonically defined areas PFt, PFop, PF, PFm, PFcm, PGa, and PGp[23]. Posterior IPL areas, PFm and PGa integrate visuo-motor, visual object, and reward input and connect with the hippocampal system, also provide access to motion-related STS areas; and PGp involves in transforming coordinates and in "idiothetic update of hippocampal visual scene representations"[24]. |
| | | Area 10r (vmPFC) | Area 10r plays a crucial role in stimulus-oriented attention and working memory (Burgess et al., 2007). |
| | | POS1 (pCunPCC) | As above |
| **22** | HIP-Head-m2 | PHC | As above |
| | | POS1 (VC) | As above |
| | | TGd | "TGd and TGv make up the temporal polar cortex, a paralimbic region important for social and emotional processing, auditory and visual aspects of facial recognition, emotional processing of auditory, olfactory and visual stimuli, and theory of mind. It also plays a role in |

| | | | |
|---|---|---|---|
| | | | language processing, especially in the ventral stream of language comprehension”[62]. |
| | | EntC | As above |
| | | STSda | As above |
| | | PGp (IPL) | As above |
| | | Area 10r (vmPFC) | As above |
| | | POS1 (pCunPCC) | As above |
| | | CER | As above |
| **23** | HIP-Head-l | PHC | As above |
| | | PerC | As above |
| | | TGd | As above |
| | | EntC | As above |
| | | STSda | As above |
| | | Area a47r (vlPFC) | As above |
| | | Area 10v (vmPFC) | Area 10v is often associated with the ventromedial prefrontal cortex (a crucial node in DMN) and involves in decision-making, complex planning, working memory, and integrating information related to reward and risk assessment from OFC and ACC(Grabenhorst & Rolls, 2011). |
| | | Area 10r (vmPFC) | As above |
| | | Area 9m (dmPFC) | As above |
| | | POS1 (pCunPCC) | As above |
| **24** | HIP-Body | PHC | As above |
| | | TGd | As above |

| | | | |
|---|---|---|---|
| | | Area 47s (vlPFC) | Area 47 involves in top-down semantic retrieval and executive control in language processing[32,33]. In addition to its connection with Broca's area, Area 47 is also considered part of the broader "Broca's complex," which also includes Brodmann areas 44, 46, 47, and the mesial supplementary motor area of 6. Together, these regions form a frontal-subcortical circuit[34]. |
| | | Area a10p (dlPFC) | As above |
| | | Area 9m (dmPFC) | As above |
| | | CER | As above |
| **25** | HIP-Tail | PHC | As above |
| | | VMV1 | VMV1 involves in the "integration of color, texture, and form information for the holistic recognition of places" [63]. |
| | | V2 | As above |
| | | POS1 | As above |
| | | Area 2 (S1) | As above |
| | | AIP | AIP involves in shaping the hand for grasping objects with the help of inputs from object recognition area in inferior temporal cortex and ventral and dorsolateral visual streams[64]. AIP also involves in tactile shape-processing and understanding orientation in space[65]. |
| | | PoI | As above |
| | | PerC | As above |
| | | PF (IPL) | As above |
| | | POS2 (pCunPCC) | As above |
| | | CER | As above |
| **26** | AMY-l | STGa | As above |

| | | | |
|---|---|---|---|
| | | PoI | As above |
| | | TGd | As above |
| | | PerC | As above |
| | | STSda | As above |
| **27** | AMY-m | Area 4 (M1) | As above |
| | | PH (TPOJ) | “Area PH is a higher-level holistic perception region of the visual system that acts as a hub of ventral stream input, integrating place-specific information. Area PH encodes, holds, and subsequently reactivates representation of a local scene, implicating it in the formation of spatial maps, place encoding, and place recognition”[66]. |
| | | Area 6r (SMA) | “Area 6r is functionally related to Broca’s area in humans, which is a well-known cortical area essential to language processing”[67,68]. |
| | | STGa | As above |
| | | Area 13l (OFC) | The orbitofrontal cortex plays a central role in processing emotions, reward value, and the experience of non-reward such as when an expected reward is not obtained[69]. In humans, the medial orbitofrontal cortex is involved in processing various types of rewards, while the lateral orbitofrontal cortex is linked to non-reward and punishment. Failing to receive an expected reward can result in feelings of sadness or depression. It is known that depression is associated with overactivity in lateral orbitofrontal cortex (which processes non-reward) and underactivity in medial orbitofrontal cortex (which processes reward). Specifically, area 13l serves as hub for distinct sensory signals delivering reward information[70] and further, integrating this information with current internal states for overall valuation of stimuli[71]. |
| | | PerC | As above |
| | | TGd | As above |
| | | Area 8BM (dmPFC) | As above |

| | | | |
|---|---|---|---|
| | | STSda | As above |
| | | A5 (STG) | Auditory Area A5 is part of the auditory association cortex and primarily responsible for processing complex acoustic sounds, including speech and environmental noises, particularly at the level of perception and concept formation. It helps interpret and understand the meaning of sounds rather than basic sound detection[72] |
| | | Area 47s (vlPFC) | As above |
| | | Area a47r (vlPFC) | As above |
| | | Area 10r (vmPFC) | As above |
| | | Area 44 (IFC) | As above |
| | | Area 9m (dmPFC) | As above |

*ROI names from Glasser atlas [73]; a – anterior, d – dorsal, l – lateral, m – medial, p – posterior, pr – prime, r – rostral, s – superior, v – ventral, A1 – Primary Auditory Cortex, A5 – Fifth Auditory Area, AIP – Anterior Intra Parietal, AVI – Anterior Ventral Insula, CER – Cerebellum, EntC – Entorhinal Cortex, FC – Frontal Cortex, FOP – Frontal Opercular, IFG – Inferior Frontal Gyrus, INS – Insula, IPL – Inferior Parietal Lobule, M1 – Primary Motor Cortex, MCC – Middle Cingulate Cortex, OFC – Orbitofrontal Cortex, OP – Parietal Operculum, PF – Parietal area F, PG – Parietal area G, pCunPCC – Precuneus-Posterior Cingulate Cortex, PerC – Perirhinal Cortex, PFC – Prefrontal Cortex, PHC – Parahippocampal Cortex, PMC – Premotor Cortex, PoI – Posterior Insula, POS – Parieto-Occipital Sulcus, PreACC – Pregenual Anterior Cingulate Cortex, S1 – Primary Somatosensory Cortex, S2 – Secondary Somatosensory Cortex, SCEF – Supplementary and Cingulate Eye Fields, SFL – Superior Frontal Language Area, SMA – Supplementary Motor Area, STG – Superior Temporal Gyrus, STS – Superior Temporal Sulcus, TPOJ – Temporo-Parieto-Occipital Junction, TG – Temporal Gyrus, V1 – Primary Visual Cortex, V2 – Second Visual Area, VC – Visual Cortex, VMV – Ventromedial Visual Area
